# A trace-based analysis pipeline for coherent and optimized electrophysiological data analysis

**DOI:** 10.1101/2025.03.25.645208

**Authors:** Julien Ballbé, Margaux Calice, Marta Gajowa, Lyle J. Graham

## Abstract

The development of large-scale neuronal networks notably relies on the use of point-neuron models to reduce the computational cost of simulations while focusing on integrative neuronal properties. However, the precise tuning of these neuron models remains a major aspect of modeling work to accurately reproduce neuronal properties and understand their implications in network activity. To this end, the precise characterization of neuronal electrophysiological properties, from linear properties to the input-output (I/O) relationship and spike frequency adaptation, from intracellular recordings is a crucial step. Furthermore, the increasing availability of publicly accessible databases opens the possibility of deriving I/O properties for point-neuron models from multiple datasets studying different neuronal populations. However, despite recent advancements in establishing universal data formats for electrophysiological studies, challenges persist due to the absence of standardized protocols (notably for current-clamp experiments) and unified data analysis methods, hindering cross-database comparisons of electrophysiological features.

To address these limitations, we developed the TACO pipeline, a free, Python-based pipeline for analyzing databases of current-clamp recordings. The TACO pipeline is designed to be user-friendly, minimizing the need for manual implementation of database-specific data extraction methods and enabling the application of user-defined quality control criteria. The pipeline incorporates robust methods for characterizing neuronal I/O relationships, spike-related feature adaptation, and estimating common experimental artifacts such as bridge errors. These methods have been designed to accommodate variability in database-specific experimental design, the sampling of the input space being of particular importancee. We validated the utility of this approach by demonstrating performance comparable to or exceeding that of machine learning models reported in the literature for neuronal type classification, using protocol-agnostic features extracted by the pipeline. This work highlights the potential of database-independent data analysis tools to enhance cross-database comparability and interoperability, advancing research sustainability and promoting the principles of Open Science.

## 2 Introduction

The study of large-scale neuronal networks constitutes a major aspect of modern computational neuroscience to decipher the underlying mechanisms of cerebral properties. Several implementations of network models had relied on multiple different approaches, benefiting from the extending computational resources available to consider fully detailed neuronal models (Markram et al., 2015; Borges et al., 2022), or a combination of fully detailed and reduced point neuron models (Billeh et al., 2020). The use of reduced point neuron models to perform large-scale network simulations (Potjans and Diesmann, 2014; Duarte and Morrison, 2019) has drawn even more attention, as they are more computationally efficient, by ignoring neuronal morphology, therefore allowing for the modeling of larger networks, even though some point neuron models have been implemented to reproduce morphologically distributed dendritic computation (Li et al., 2019), and lead to the investigation of neuronal functional properties in network dynamics.

### 2.1 Definition and biological relevance of neuronal firing characteristics

Neurons are the building blocks of the nervous system and represent the elementary computational unit of cerebral networks, as all neurons in a network receive signals coming from the rest of the network, integrate them and generate a corresponding response which is sent downstream. The way in which a neuron transforms the received inputs into a reproducible output, and how this Input-Output (I/O) relationship can be described and modulated constitute a wide field of studies. Indeed, different metrics can be derived from this relationship, each representing a crucial aspect of neuronal biophysics (Silver, 2010).

The rheobase is defined as the minimum input needed to produce an action potential. This property inversely describes neuronal excitability (i.e.: increased neural excitability comes with a decrease of neuron’s rheobase) and can be modulated through diverse mechanisms like Ca^2+^ release from endoplasmic reticulum or reduction of Ca^2+^-sensitive potassium current (Shine et al., 2021). Rheobase has been reported to play important role in network properties. Notably, distance to threshold heterogeneity has been observed to be reduced in epileptogenic tissues, leading to disrupted relationship between driving input and network synchrony when rheobase variability was limited in excitatory neurons (Rich et al., 2022). Additionally, inhibitory neurons’ rheobase variability was reported to make network’s asynchronous state more robust (Mejias and Longtin, 2014).

The gain is defined as the slope of the I/O relationship and describes the neuron’s output sensitivity to a change in input (Salinas and Sejnowki, 2001). As for rheobase, gain is not fixed and can be modulated through different mechanisms like input correlation (Salinas and Sejnowski, 2000), or through cortical inhibition(Ferguson and Cardin, 2020). At the network level, neuronal gain has been reported to play important role on higher cortical functions. Indeed, gain’s variability has been observed to encode for stimulus uncertainty, notably in the context of visual stimulus (Hénaff et al., 2020; Festa et al., 2021). While rheobase and gain are commonly investigated neuronal I/O features, other aspects of this relationship can be measured like the saturation stimuli corresponding to the stimuli eliciting the neuron’s maximum response, and over which the response of the neuron either saturates or decreases.

### 2.2 Complex description of neuronal I/O relationship and firing properties

Each of these measures allows to characterize the neuron’s response to stimulus and can play important roles in network’s behavior and properties. Therefore, carefully characterizing neuronal response represents a crucial aspect of neuronal physiology and computational properties, yet no consensus exists as regard to its description and characterization.

One main reason explaining this lack of universal method, is notably because of context-dependent nature of the I/O relationship. Indeed, while in the context of a current-clamp experiment, a neuron receives a defined current impulse for a defined duration and emits trains of action potentials, in conductance-clamp, the input can be composed of two different conductance components, one excitatory and one inhibitory. Furthermore, in the context of *in-vivo* experiment characterizing the neuronal response to visual stimuli, the input can be considered as the orientation of the visual cue presented to the animal (Hubel and Wiesel, 1962). In the context of current-clamp experiments, the input is represented by the amplitude of the current applied to the cell, generally expressed in pico-amperes (pA), while the output of the neuron is represented by the train of action potentials emitted in response to this input. While the definition of the different firing properties (i.e.: rheobase, gain, …) are globally common, the methodology employed to describe the I/O relationship may considerably vary between different studies. The common guidelines to characterize the I/O relationship from a current-clamp experiment consist in applying current steps of increasing amplitude, record the corresponding change in membrane potential, detect action potential over the stimulus duration, and get the firing frequency corresponding to the input level. A function is then generally fitted to the input-frequency curve (f-I) to obtain a more stable estimation of the rheobase, gain, and eventually of the saturation. However, several experimental aspects may limit the comparison of I/O data gathered in different experiments.

First, the experiment consists in making a discrete sampling of a continuous space (i.e.: a neuron can receive infinite different levels of input). Therefore, the more sampling of the I/O space, the more precise will be the characterization of the I/O relationship, but that requires longer recording sessions during which the stability of the patch-clamp or the cell’s health can deteriorate. Thus, it is common to sample the input space with wider current steps to ensure entering the response regime of the neuron, but therefore reducing the resolution of the sampling, and the specificities the I/O relationship may have. Furthermore, no consensus exists as to what is the optimal current steps amplitude, adding a technical source of variability between different databases.

Secondly, the fitting procedure plays a crucial role in the subsequent estimation of firing properties, even though two cells may have been sampled similarly. Indeed, while the definition of the I/O features we saw before are generally shared, different functions can be used to fit the I/O relationship, like simple linear fit (Fernandez et al., 2015; Gouwens et al., 2019), polynomial (Chance et al., 2002), logarithmic (Mitchell and Silver, 2003), or sigmoid function (Fellous et al., 2003; Cardin et al., 2008) (see Figure 1). The use of different fitting functions to describe similar I/O relationship can lead to significant different conclusions as to the general shape of the transfer function. Except in the case of linear fit, all other examples allow to characterize crucial aspects of the I/O relationship: the smooth transition between silent state of the neuron and the linear phase of the I/O relationship may vary in shape, being either convex or concave, as well as spotting potential saturation of firing frequency, but not potential frequency decrease after the saturation stimuli.

**Figure 1:**
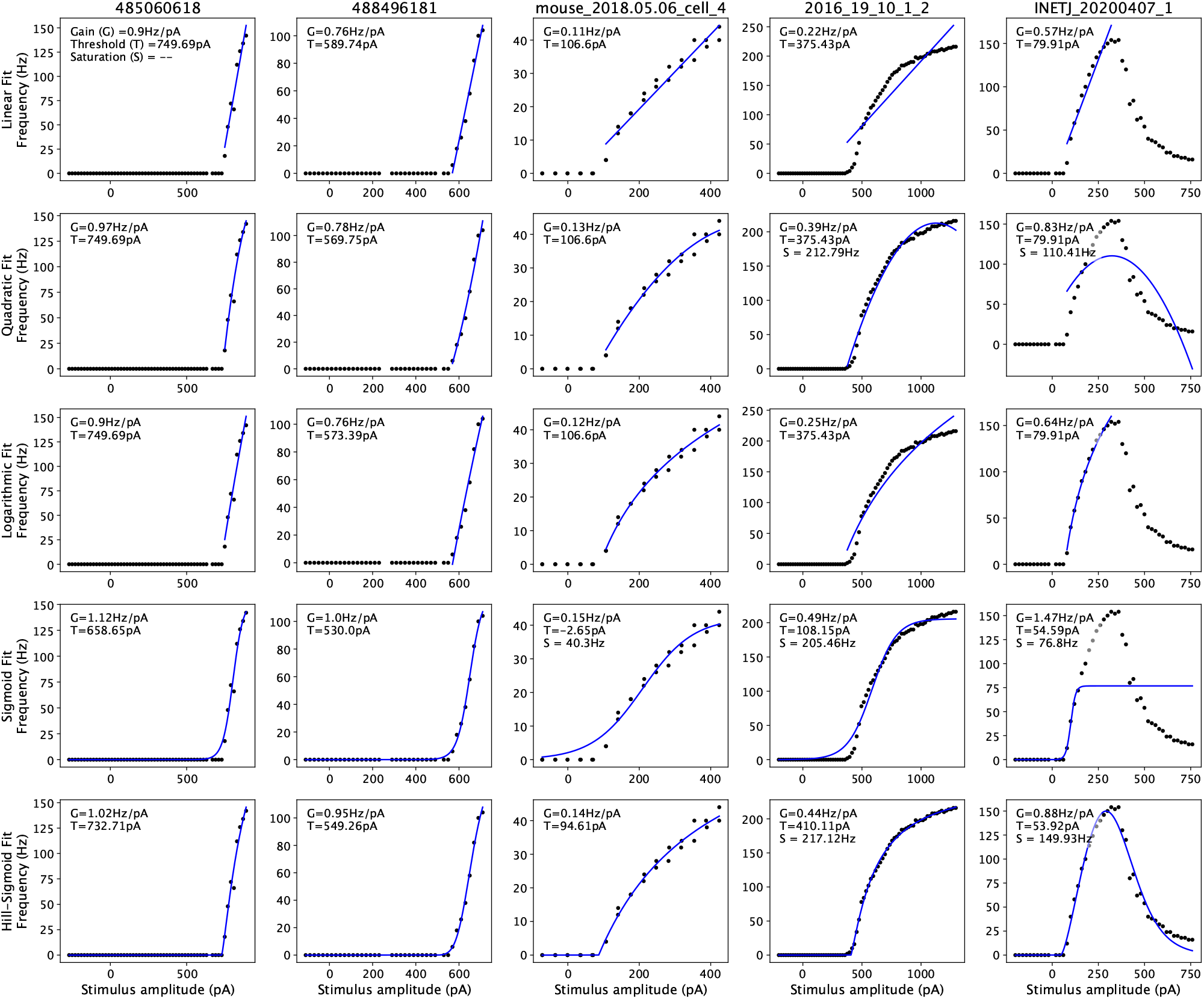
Comparison of different IO fitting procedures found in the literature and proposed by the laboratory (i.e.: Hill-Sigmoid) Neurons display a wide diversity of electrophysiological properties, some directly related to the IO relationship like the Gain, the Threshold or the Saturation. Yet the choice of fitting function has a direct influence on the estimation of those features. Fitting functions were chosen in the literature, linear (Fernandez et al., 2015; Gouwens et al., 2019), second order quadratic (Chance et al., 2002), logarithmic (Mitchell and Silver, 2003), or sigmoid function (Fellous et al., 2003; Cardin et al., 2008), and implemented as specified in the references or inferred from the figures in the references. The Hill-Sigmoid fit has been developed in the laboratory to account for the diversity of IO relationships. In each case, the gain was calculated as a linear regression to the mid portion (mid 50%) of the fit. Threshold was determined as the minimum value of the fit corresponding to the minimum observable response (i.e.: the response duration considered being 500ms, the minimum observable was one action potential so 2Hz). The saturation was detected if the ratio of the final and maximum slope of the I/O fit was less than 0.2. If so, the saturation frequency was considered as the maximum value of the I/O fit, and the saturation stimulus as the corresponding input. Cells were taken from Allen Cell Type Database (485060618, 488496181), NVC database (mouse_2018.05.06_cell_4), Scala 2019 database (2016_19_10_1_2) and Scala 2021 database (INETJ_20200407_1)

Finally, apart from the methodological consideration we described, a fundamental biological aspect of neuronal firing can render a general I/O fitting procedure harder to implement. In 1948, Hodgkin described three fundamentally different type of neuronal firing, commonly referred to as Type I, Type II and Type III neurons (Hodgkin, 1948), each having distinct characteristics. Type I neurons, also be called integrators, are capable of repetitive firing over a wide range of frequencies, even low range frequency (some Hz), making them ideally suited to encode stimuli intensity. Type II neurons (also called resonator) present the particularity to firing only in high frequency range (at least several tens of Hz), and without particular dependence on the input strength. Finally, Type III neurons gather cells unable to sustain repetitive firing, which either fails or are limited to firing a single action potential even for high level stimuli (Prescott et al., 2008a). While the concept of I/O relationship hardly applies to the late class, it is different between the first two classes, implying fundamentally different spike firing dynamics. Indeed, Type I neurons’ I/O relationship (while all different) are presented as continuous while on the other hand, Type II neurons can only fire action potentials at high rates and are unable to fire spikes at low firing rates, making their I/O curves looking like a discontinuous relationship because of the big frequency step between the subthreshold and firing regimes (Izhikevich, 2007; Prescott et al., 2008a). If the computational implication of these different excitability classes has been well studied (Zeldenrust et al., 2021), the boundary between these types does not seem to be an absolute classification as neurons have been observed to shift between the two classes under different conditions (Prescott et al., 2008b; Zhao and Gu, 2017). This shifting may explain why, despite the clear qualitative difference between Type I and Type II neurons, that is the continuous and discontinuous aspects of their I/O curves, there are no clear quantitative definition as to what makes an I/O relationship of Type I or Type II. Indeed, when performing current-clamp experiment, and so sampling the input space, identifying a discontinuity in a discontinuous space becomes tedious. Some works have been done on trying to quantify a minimum firing frequency under which Type II neurons cannot sustain repetitive firing, estimating a lower-bound between 10 and 30Hz (Tateno et al., 2004) or between 20 and 30Hz (Tateno and Robinson, 2006). Yet such frequency steps are not uncommon, especially in experiments with low input resolution (i.e.: with big current amplitude steps), making the identification of a clear boundary between the two categories difficult. Therefore, the application of a single fitting procedure to all neurons’ I/O relationship is a delicate subject, as one should define from current-clamp recording what makes a neuron Type I or Type II.

### 2.3 Spike frequency adaptation, an ubiquitous property

Another biophysical neuronal biophysical property is the progressive decrease of firing frequency, generally known as spike frequency adaptation (SFA), which is a crucial aspect of a neuron’s response to external stimuli. SFA is commonly defined as the evolution of instantaneous spike frequency over a sustained input received by the neuron and seems to be an ubiquitous neuronal property (Fuhrmann et al., 2002; Ha and Cheong, 2017), although expressed in different degrees in neuronal population. This property has notably been used to classify neuronal types (Ascoli et al., 2008). The processes giving rise to this phenomenon are multiple and involve recent spiking activity-dependent mechanisms. These mechanisms can involve voltage-dependent activation of M-type currents, or after hyperpolarization current activation in response to increased calcium concentration (Benda and Herz, 2003; Gutkin and Zeldenrust, 2014; Ha and Cheong, 2017). The effects of these mechanisms have been reproduced computationally in neuronal models, notably in integrate-and-fire derived models, by adding after-spike currents, after-spike AHP dynamics, or by adding dynamic spike threshold, even though these different mechanisms may induce subtile differences in the model behavior (Benda et al., 2010). In the perspective of its biological relevance, SFA has been observed to play important role in neuronal network properties. It notably has been reported to provide network with better temporal and working memory capacities (Salaj et al., 2021), and extend stability regions for chaotic neural networks (Muscinelli et al., 2019).

Regarding SFA quantification, the general method consists in considering the inverse of consecutive inter spike interval (ISI) (Mitrić et al., 2019) or instantaneous frequency as an exponentially decaying value over a time period (i.e.: the input duration), making SFA a time-based mechanism. While this represents a convenient and reliable method, as for I/O fit, such definition lacks precision to be universally applied on multiple heterogeneous data, especially for current-clamp experiments where they can be prone to over-fitting, and vary because of the protocol discrepancies between databases. Attempts at defining more robust method have been made, notably by considering two different measures of ISI variation, one local and one global over a spike train (respectively Lv and Cv). Such quantification method in spike trains recorded during various tasks appeared to be able to classify neurons based on these measures (Shinomoto et al., 2003, 2009).

Therefore, the quantification of neuronal biophysical properties is a practical matter, often addressed individually by laboratories, in the context of their own database. Yet, such approach can prevent from coherent analysis comparison between databases, as one laboratory’s measure may not be appropriate to other laboratory’s experimental protocol. Thus, the development of methods to characterize the intrinsic biological diversity of neuronal I/O relationship and firing properties such as SFA, while reducing experimentally-induced variability represents a key subject of computational neurosciences.

To address this question, we implemented a set of data-driven methods to quantify functional neuronal properties, ensuring consistent evaluation of these properties from differently implemented current-clamp recordings. To enable better interoperability between neuronal properties measured from different databases, these methods are implemented as part of a Trace-based Analysis pipeline for Coherent and Organized database organization (TACO pipeline), a Python-based semi-automatic data treatment pipeline. We use this pipeline on five publicly available databases (Harrison et al., 2015; da Silva Lantyer et al., 2018; Gouwens et al., 2019; Scala et al., 2019, 2021), as well as on the database implemented by the laboratory. Our work represents a step toward the implementation of methods more robust to differences in experimental protocol, enabling more coherent analysis across databases. Furthermore, the TACO pipeline has been designed to be applied in a variety of database’s structures, compensating for the differences in curation between databases, and enabling the application of user-defined quality criteria, ensuring coherent and consistent comparison between databases. Such design notably aligns with the guidelines of Open Science, promoting better interoperability between data and inter-databases comparison.

## 3 Trace-based Analysis pipeline for Coherent and Organized databases integration (TACO pipeline)

There are non-negligible number of publicly available databases of intracellular recording which are worth studying and integrating. Yet only few of them follow common data format such as NWB data format (Rübel et al., 2022). To overcome this and propose an alternative solution toward open neurosciences, we designed a python-based data-treatment pipeline, with the aim of valorizing existing databases of intracellular recordings. This pipeline has been designed based on the hypothesis that three main aspects of currently available databases limit their integration altogether.

1. Databases describing experiment of intracellular electrophysiological recording, (e.g.: current clamp experiment) gather similar type of raw data (i.e. current and voltage traces). Yet, most of them are acquired, organized, stored and analyzed in a lab-fashion way. Therefore, it often represents a significant amount of time to understand the particular organization of the database, before being able to rely on the analysis that has been made on it or re-organizing it so it can undergo custom analysis.
2. While the data may be acquired with similar original aim, the particular implementation of the protocol may vary between laboratories and even within a each of them. For example, in a current-clamp experiment where the variation of cell’s membrane voltage resulting from the injection of different levels of current is recorded, several aspects can be tuned differently by different labs, like the amplitude or the duration of the stimuli, or the number of stimuli applied. These small, yet significant, differences between databases make the use of a common analysis pipeline able to accurately characterize the various aspects of the data complicated.
3. Using different databases alongside with each other represents both a purely scientific interest and addresses the concern of science sustainability, as maximizing existing data represents key aspects of modern open neurosciences, notably to limit animal experiments Janssens et al. (2023); Richter (2024). Yet gathering similar data generated by different labs, in different conditions, with different purposes, implies to keep a critic eye, to be sure not to compare apples and oranges (Purgato and Adams, 2012). Indeed, while data will inevitably present endogenous heterogeneity, it is crucial to be able to identify extra-sources of heterogeneity, like animal to animal variability (i.e.: individual experience) or the experimental conditions (e.g.: temperature, materials, pre-processing, …) (Pastoll et al., 2020).

Therefore, the pipeline has been designed as a general, participative framework for databases integration and analysis, where each lab could participate to the integration of their database, while assessing quality statements they would deem necessary to compare other databases to their own (Figure 2).

**Figure 2:**
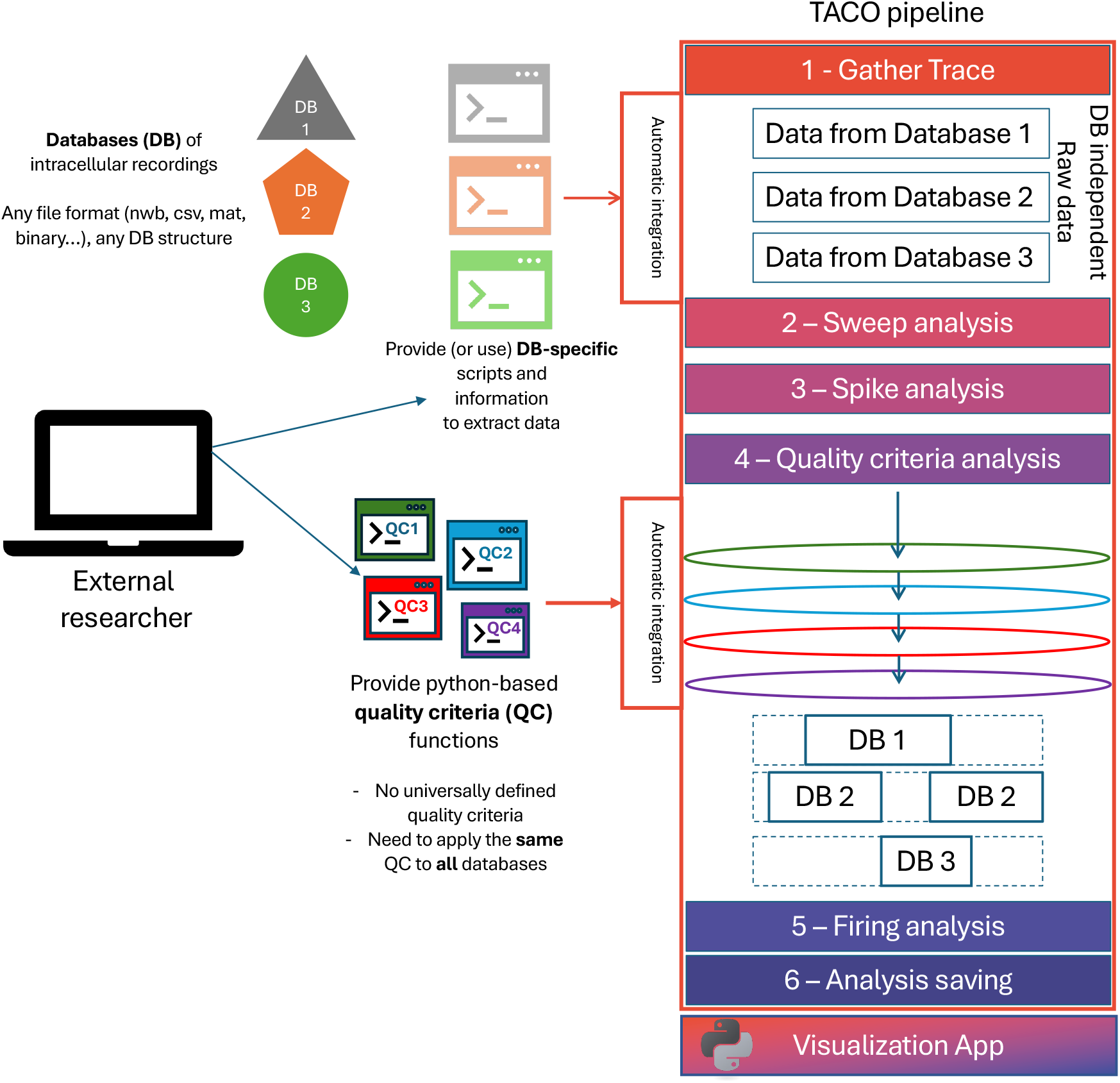
TACO pipeline organization: The TACO pipeline is designed to integrate and analyze databases of intracellular recordings independently from their structures and file format. **1 - Gather Trace** Based on the database-relative information and data extraction scripts provided by the user and automatically integrated by the pipeline, the raw traces of the databases are extracted and organized for the analysis. **2 - Sweep analysis**: for a single cell, the traces are analyzed to extract relevant sweep-related data like stimulus times and amplitude, estimate bridge error, and compute sweep-based neuronal linear properties like resting potential, input resistance or time constant. **3 - Spike analysis**: for each sweep, the potential trace is analyzed to extract spike-related information like time and potential value of threshold, peak, trough or maximum rate for the increase and decreasing action potential phase (i.e., upstroke and downstroke). This analysis is based on the analysis provided as part of the AllenSDK (Gouwens et al., 2019). **4 - Quality criteria analysis**: To ensure consistency across databases, the user of the pipeline can provide a set of python-based functions describing sweep-based quality criteria (QC). This step has been designed so that the user can directly refer to sweep and cell-related properties (current trace, potential trace, input resistance, bridge error, …), and the different QC will be assessed automatically without any need to manually implement these into the pipeline. Any sweep failing any of the QC proposed by the user will not be considered for the firing analysis, or the cell-based computation of linear properties (solid rectangles represent the kept traces for each database). **5 - Firing analysis**: The QC-validated sweeps are then used to compute the firing properties of the neuron, by performing data-based fit of the input-output (I/O) relationship. **6 - Analysis saving**: The different aspects of the analysis are saved in an HDF5 file format to enable further study of the analysis, and the visualization of the analysis using the python-based visualization app.

### 3.1 Pre-requisite: Defining common terms

Before digging into the details of the pipeline, it is important to start by giving a set of definitions used in the context of the pipeline and referring to different aspect of the current-clamp experiment. Notably we define a “trace” as an array of recorded values representing a unique modality of the experiment (i.e. voltage trace, or current trace). A “sweep”, is a set of corresponding voltage and current traces for a given cell. A sweep is referred to by a single identifier “sweep id” which is unique for a given cell but can be used for different cell. A “protocol” is constituted by a train of sweeps grouped together as they are done one after the other and have unique level of stimulation. Finally, an “experiment” refers to the ensemble of the protocols that have been performed on the cell. (See Figure 3).

**Figure 3:**
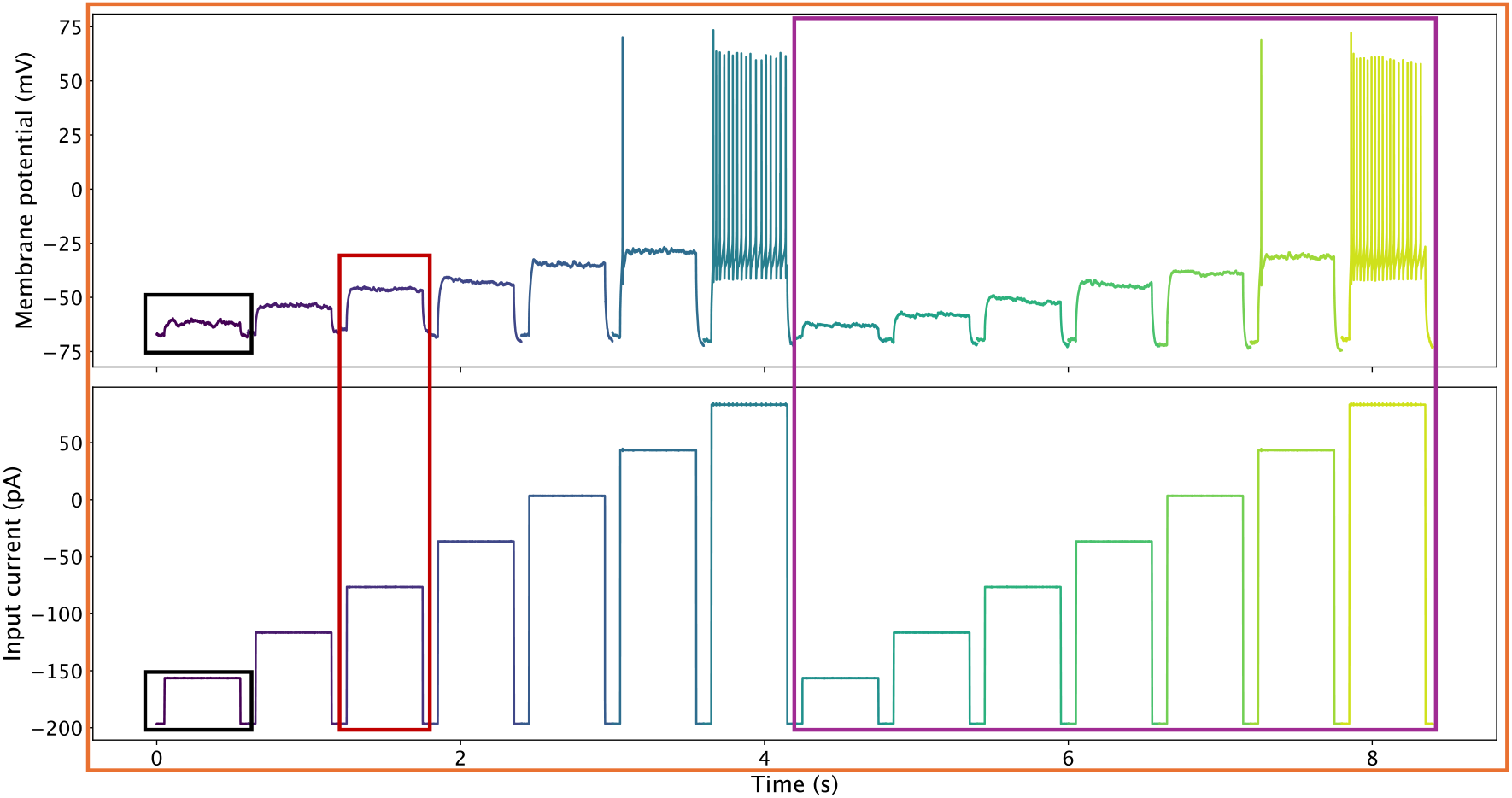
Definition of the different components of an electrophysiological recording in current clamp configuration. A trace is the voltage or current array of recorded value (black rectangles). The combination of corresponding voltage and current trace constitute a sweep (red rectangle). A protocol (purple rectangle) corresponds to the ensemble of sweeps performed one after the other, with unique an increasing level of stimulation. Finally, an experiment (orange rectangle) corresponds to all the protocols performed on a cell

### 3.2 User friendly environment and required information

#### 3.2.1 Python as an easy to use framework

Modern neurosciences research heavily relies on computational tools for data analysis. Different scientific-oriented coding language exist, each having their own specificities, advantages and drawbacks. For example, MATLAB is a programming platform used in various application from data processing to machine learning, or image processing. Furthermore, it proposes Simulink, a graphical interface to design processing pipelines. While powerful and versatile, this is a commercial product, therefore not open source, making it a less accessible for researchers not interested in more advanced features. R is an open-source programming platform, mainly used in the context of statistical analysis and graphical data investigation, relying on the use of different processing libraries. While it does not provide a graphical user interface natively, different graphical libraries have been developed through the years, like GGplot2 to generate plots, or Shiny to develop graphical application relying on R code for data processing. Finally, Python is an open source programming language and, like R, a procedural, object-oriented framework. It too relies on external libraries to integrate more processing capacities, making it a versatile solution for laboratories. Furthermore, its human-readable syntax makes it an easy to learn programming environment, and a self-sufficient solution for laboratories as it provides solution for various scientific tasks, from data processing, statistical analysis, pipeline implementation, web management, and even machine learning. Its versatility progressively imposed it as an essential tool, notably in the neuroscience community, as it has been used in various large-size projects like PyNN (Davison et al., 2009), PyNEST (Eppler et al., 2009) or BMTK (Dai et al., 2020) to say a few. Therefore, as the use of Python is widespread in neurosciences research, most researcher have at least basic experience of coding using function definition, which makes it a good option for a common data treatment pipeline.

#### 3.2.2 Requirements for database integration

As stated before, each lab could participate into the integration of their database. To do so, the key element of participation resides on the basic access to raw data, and the listing of recorded traces present in the database. More precisely, to integrate any database of current-clamp recordings in the pipeline, a lab only needs to provide some common information about the current-clamp experiment.

1. **Population class table**: The first document required is a csv file specifying and describing each cell present in the database. For example, studies generally focus on specific neuronal cell types, on particular cortical areas or specific layer. Apart from the cell id and the name of the database it belongs to, no information is mandatory. However, the more details can be attributed to each cell, the more accurate will be the analysis performed on the database, and more consistent will be the comparison of data coming from different databases. Such data can relate to cellular information (e.g.: cell type, custom classification, cortical area, …) or to recording conditions (e.g.: age of the animal, recording temperature…)
2. **Cell-Sweep table**: The second element required by the pipeline, is a csv file describing, for each cell present in the database, the sweep to consider for the analysis. This information mainly concerns the organization of the database. The idea is that in any lab, any experiment performed on a cell is fundamentally a collection of sweeps. Most of the time, databases of current-clamp recordings organize their data so that different sweeps can easily be accessed, notably by indexing them separately by giving them a unique sweep id. However, it should be noted that some databases may not directly provide such information directly but rather provide method to understand the storage of the experiment, which results in the a-posteriori definition of sweep ids. Also, some databases may store in a same file, multiple kind of recordings. This is notably the case for the Allen Cell Type Database(Gouwens et al., 2019) which recorded for each cell different protocols of current-clamp experiments (i.e.: long square stimulus, short square stimulus, noise…). By specifying this information for any cell of the database, it allows to ensure only the same kind of protocols are considered, to keep track of the sweeps present in an experiment and organize the analysis in a trace-based fashion.
3. **Trace extraction Python script**: Finally, the last required element for the integration of a database, is a Python code defining a function, which given a path toward a cell file, and the list of sweep id of interest, returns the corresponding voltage, current and time traces. This requirement is at the core of gathering different databases together, as it is the only portion of the pipeline relative to the structure of the original databases. This function only describes how to access the raw data for a given database, without any need to manually integrate it into the pipeline. Indeed, the pipeline automatically import the function, provide it with relevant inputs, and receives the corresponding traces which will undergo the analysis. If the database contains the stimulus start and end times, the function can also return them. However, if it doesn’t, the pipeline will automatically estimate the stimulus start and end times by performing autocorrelation on the current trace first time derivative, based on the stimulus duration provided by the database. The advantage of this approach is to remove the barrier of database’s structure by extracting the raw data and feeding them into the analysis pipeline, without any need of pre-treatment, tuning, or restructuration of the database. Therefore, we separated the data extraction from the actual analysis. Yet a strong source of variability can still be present in the data and is the direct consequence of having different data acquired by different experimentalists, using different tools. Therefore, such kind of experimental artifacts should be quantified and/or compensated, to ensure the subsequent analysis, and the comparison of different databases is reliable.

### 3.3 Ensuring reliable analysis for cross-comparison between databases

#### 3.3.1 Bridge balance correction, a common artifact in electrophysiological recordings

Recording the dynamics of the membrane potential is a key aspect to characterize the electrophysiological properties of a neuron. Indeed, the neuronal membrane is composed of a bi-lipid membrane, creating an insulating layer between two conducting electrochemically different solutions (i.e., intra vs extra-cellular media), giving to the membrane the properties of a capacitor, maintaining a difference of electronic charges across the membrane. Yet the membrane also contains numerous trans-membrane ionic channels assuring the passive or active transport of ionic molecules across the membrane. Each of these trans-membrane channels represents a particular conductance (or resistance), making variable the membrane potential upon the application of an external current. Collectively, these channels determine the membrane’s *Input Resistance*, a key indicator of the cell’s excitability typically measured when the cell’s is at its resting potential. Input resistance therefore reflects how much the membrane potential varies in response to an input current. However, when performing patch-clamp experiment the cell is sealed to a glass electrode linked to the amplifier. From this experiment, different configurations can be implemented depending on the question of interest, and enabling the electrical access to the inside of the cell.

##### 3.3.1.1 Bridge error in intracellular recordings

Patch-clamp can for example be used during current-clamp experiments in whole-cell configuration, providing electrical access to the inside of the cell, to inject a current and observe the resulting change in membrane potential. As illustrated in Figure 4, once the seal has been made between the cell and the glass pipette, a current can be injected into the cell (for the case of a current-clamp experiment), and the resulting variation of the membrane potential *V*_*m*_ can be recorded by the electrode as *V*_*p*_. However, the electrode may constitute a resistance to the flow of current toward the cell, due to various aspects like its tip size and shape, or the recording solution it has been filled up with. All these aspects constitute the *Access resistance R*_*e*_ which from an electrical point of view, is in series with the input resistance of the cell introducing an experimental artifact. Therefore, the amplifier records a potential which is the sum of the actual membrane potential (*V*_*m*_) and of the voltage resulting from the current flowing through the access resistance, or:

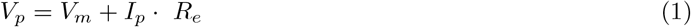

**Figure 4:**
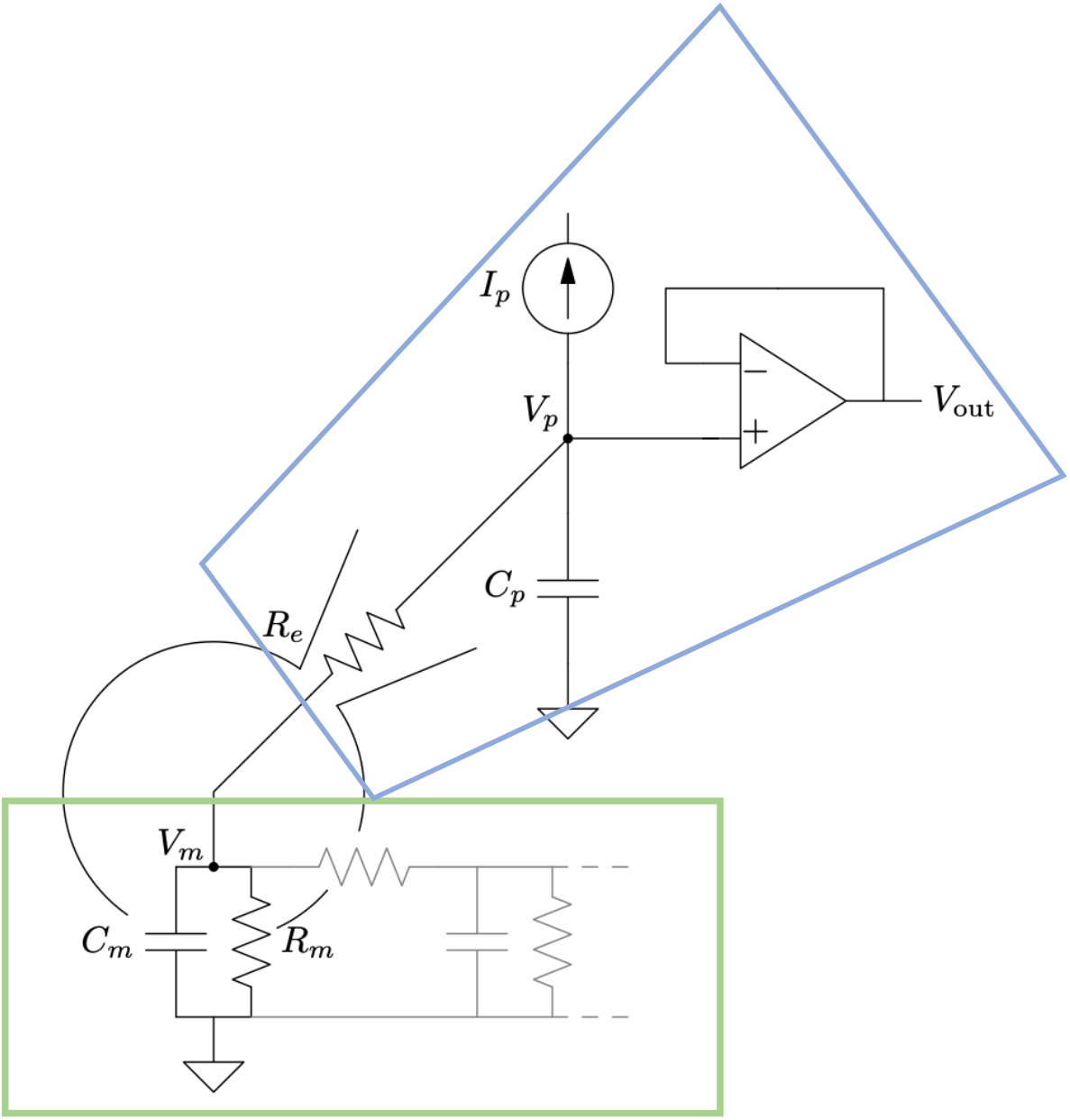
Electronic scheme of patch-clamp experiment. During a patch-clamp experiment, the electrode gets electric access to the inside of the cell by sealing the glass pipette to the cell membrane. Here, the electrode (blue shape) injects a current *I*_*p*_ into the cell (green rectangle), while the resulting variation of membrane potential *V*_*m*_ are recorded as *V*_*p*_. The fact that the access resistance (*R*_*e*_) is now in series with the input resistance of the cell (*R*_*m*_), implies the fact that the recorded membrane potential *V*_*p*_ is the sum of the actual membrane potential *V*_*m*_ with the voltage across the access resistance. Thus, it appears from this circuit diagram that *V*_*p*_ = *V*_*m*_ + *I*_*p*_ * *R*_*e*_. Adapted from figure 4 in Barbour (2014)

Furthermore, following the Ohm’s law, we can approximate *V*_*m*_ ≈ *I*_*p*_ · *R*_*m*_, with *R*_*m*_ the cell’s resting input resistance. Thus we can re-write equation 1 as:

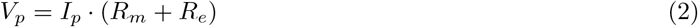

It therefore appears that at steady state (*t* → ∞) the voltage recorded by the amplifier *V*_*p*_ in response to the application of a current *I*_*p*_ is proportional to the sum of both the cell’s input resistance *R*_*m*_ and the electrode’s access resistance *R*_*e*_. Even though the electrode’s access resistance can be minimized by optimizing several factors like the tip size and diameter of the glass pipette or by using a high-conductance filling solution, it still induces undesired voltage deflection, commonly referred to as “Bridge Error”. This artifact of intracellular recordings is well known to experimentalists and is generally linearly compensated for during recording using built-in controls from the amplifier.

##### 3.3.1.2 An artifact not always compensated, the example of Lantyer Database

However, while most databases of recordings are corrected during the experiment, making the analysis of the data within the database coherent together, this aspect of bridge balance compensation becomes central when analyzing together databases acquired by different experimentalists, using different recording equipments. Also, it may happen that some databases are made of uncorrected potential traces, and do not provided information regarding the level of bridge balance compensation applied on the cell during the experiment. This is notably the case of the Lantyer database (da Silva Lantyer et al., 2018).

This work aimed at constituting a database of electrophysiological recordings that would help to characterize the electrophysiological properties of the L2/3 interneurons in the somatosensory cortex. To do so, they targeted cells in mouse lines expressing Cre recombinase for either PValb, or Sst neurons, and applied various sets of protocols, like current-clamp step and hold stimulation, triangular voltage-clamp sweeps or frozen-noise current injections. They therefore provided a rich database of electrophysiological recordings, notably containing 257 cells recorded under current-clamp configuration, making it a valuable source of neurophysiological data. However, numerous different experimentalists worked to acquire these cells, resulting in a variety of quality for the different recordings, notably regarding the bridge balance (see figure 5). Indeed, as we saw before, such uncompensated bridge balance in the raw traces can cause a mis-estimation of the actual membrane potential, and therefore lead to incorrect quantification of neuronal properties like input resistance, capacitance, resting potential, or action potential related features.

**Figure 5:**
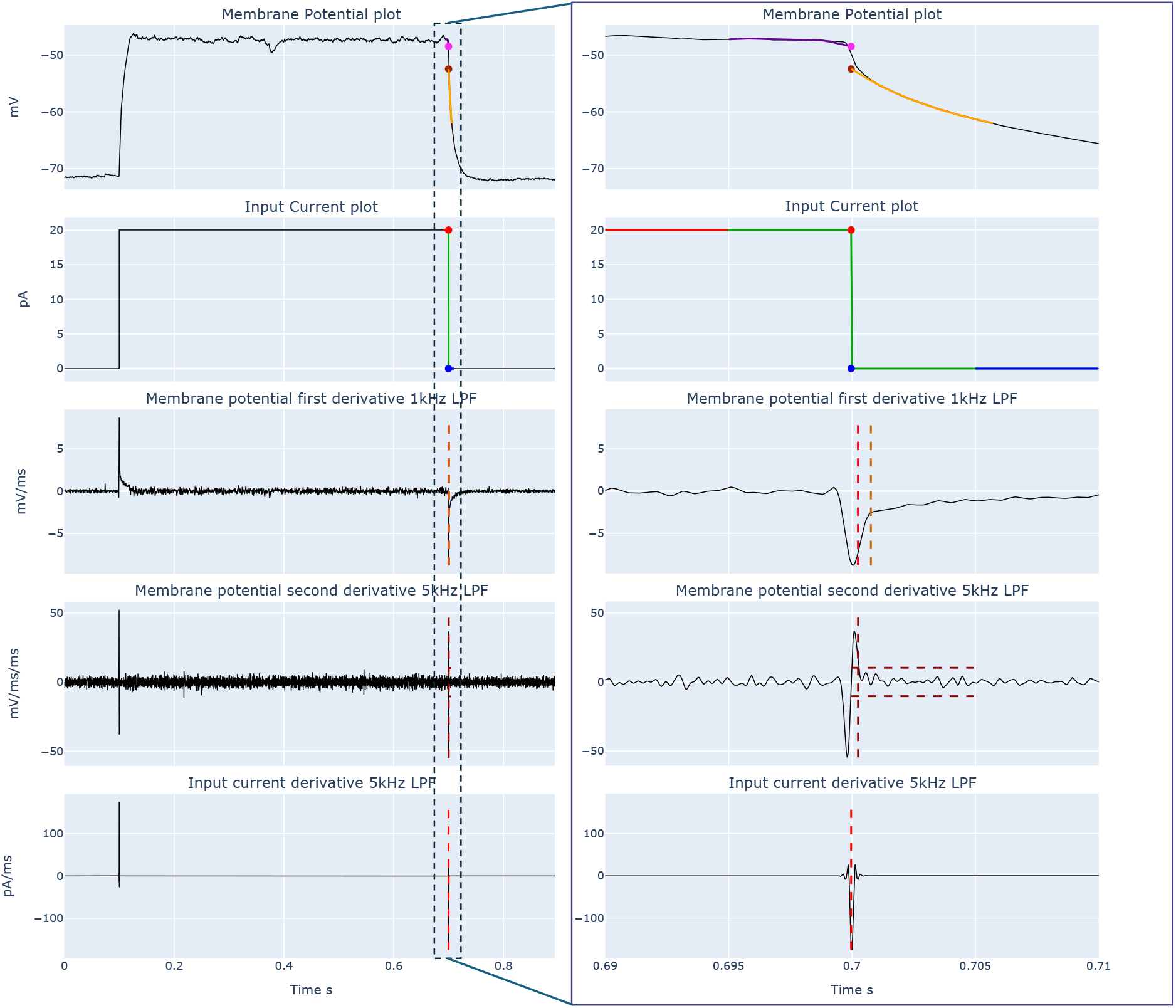
Ad-hoc bridge error compensation as presented in the cell visualization app. **Left panel**: Detailed procedure of bridge error estimation. **Right panel**: Same as left panel, zoomed in around the times of interest. **First row**: membrane potential trace, presenting the pre and post transition voltage estimates (pink and brown dot respectively), estimated as the voltage value at the time of transition from pre and post transition voltage fit (purple and orange line respectively). **Second row**: Input current trace, presenting the pre and post transition current estimates (red and blue dot respectively), estimated as the first and last current value at beginning and the end (red, and blue line respectively) of sigmoid fit on the current trace (green line). **Third row**: Membrane potential first time derivative. Red dashed line: time of reference for the start of the “true cell response” (i.e., without potential fast transient response). Orange dashed line: Start time of the membrane trace fit window (orange line in the first row). **Fourth row**: Membrane potential second time derivative. Vertical brown dashed line: Reference time for the detection of fast transient response, which are detected if the second time derivative becomes higher (or lower) than a threshold value (horizontal brown dashed lines). **Fifth row**: Input current first time derivative. The transition time is estimated as the time of the maximum absolute value of the input current first time derivative (red dashed line) In this example, the voltage drop was −3.99 mV, while the current step was −20 pA, resulting in a bridge error of 199 MΩ. This example is taken from sweep CC_5_12, cell 2017_10_05_2_6 from Lantyer Database.

It therefore represents a key element to consider, in order to assure a reliable analysis of the database, and coherent comparison with external databases.

##### 3.3.1.3 An off-line estimation of bridge error

As we saw before, the bridge error results from the addition of the input resistance of the cell with the access resistance induced by the use of the glass pipette to inject a current and record the membrane potential. It is therefore crucial carefully estimate this artifact from intracellular recordings (either current-clamp or dynamic clamp) as an experimentally induced voltage drop can highly influence the analysis made from the recording. Generally, the compensation for the bridge error is made on-line by the experimentalist. Other off-line methods relied on the estimation of the rapid voltage deflection following the stimulus transition time to estimate the bridge error, either by direct fitting to electrode-cell models (Sa and Mackay, 2001) or by kernel estimation (Brette et al., 2008). Fundamentally, the challenge resides in the distinction of the electrode-induced voltage deflection from cell membrane voltage deflection, both of which resulting from cable structures.

We therefore introduce an *ad-hoc* procedure to distinguish between the fastest time component of the cell voltage drop (i.e.: actually reflecting the input current impact on the cell membrane potential) from the electrode time constant (see Figure 5 First row). Therefore, the idea was to target the transitions time corresponding to the rapid change in input current (e.g.: at the end of the stimulus see Figure 5 Fifth row), estimate the membrane potential right before it, and estimate the membrane potential variation governed by the cell’s time constant after this transition.

The original acquired voltage and current traces from each trial were filtered with a 4th order zero-phase low pass Butterworth filter with cutoff frequency *f*_*c*_ =10kHz, to obtain *V*_*trace*_ and *I*_*trace*_, respectively. In general 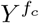 refers to a low pass filtered trace *Y* (*t*) (e.g. *V* (*t*) or *I*(*t*)) with a cutoff frequency *f*_*c*_ (Hz; 0 phase 4th order butterworth), and with 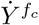 and 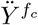 denoting the first and second derivatives of 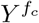.

Consider a pair of voltage *V* and current *I* traces, where *I* includes a defined sharp transition (e.g. stimulus start and stop for current clamp step protocols, or stimulus end for dynamic clamp step protocols). In general, the challenge of bridge compensation - whether done on-line by the experimentalist or off-line - is to estimate the rapid voltage jump at the stimulus transition due to the bridge error, while both taking into account fast time constants of the cell response and discounting other measurement artifacts, including any phase delay of the amplifier bridge compensation, ringing, etc.

Fundamentally the challenge is to distinguish the electrode versus cell component, both of which derive (for the linear response) from cable structures, notably with distributed capacitance. For intracellular recordings made with sharp microelectrodes, a reliable estimate of the electrode’s properties may be typically derived before entering a cell. For patch recordings, however, the electrode (“access”) resistance is highly dependent on the eventual whole-cell access configuration.

The time-domain method described here essentially formalizes the standard “by-eye” on-line procedure, with two major assumptions. The first assumption is that the upper bound of the electrode time constant is on the order of 0.5 msec. The second considers that the step response to a linear some-dendrite circuit includes a range of time constants up to the dominant intrinsic membrane time constant (Jack Nobel Tsien). Here the focus is distinguishing the faster components of the cell response from the electrode component; for that reason we assume that only the first 5 msecs of the distinguishable cell response, that is after any stimulus artifact, is relevant. All other constants, e.g. for various reference times (in msec), were set empirically.

We first check if there there are no spikes within ±25 msec of the specified stimulus transition time *T*, and if so the analysis is abandoned. To improve the accuracy of the analysis, an estimate of the actual stimulus transition time, *T* ^*^, is derived from the recorded current trace as the maximum (minimum) of *İ*^5*k*^ for a defined positive (negative) current step, within a 4 msec window centered on *T*. Measurement of the current step Δ*I* delivered to the cell takes into account the median values of *I*(*t*) in two 1 msec windows on either side of *T* ^*^ to avoid any transition artifacts of the recorded stimulus (Figure 5 Fifth row):

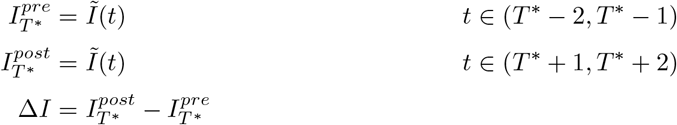

The very short windows for estimating the current step reflects the focus on the response’s fast dynamics.

Any fast ringing transient (*FT*) in *V* during 5 msecs following the stimulus transition time *T* ^*^ is detected if and when the absolute value of the voltage second derivative 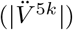 exceeds a threshold *α*_*F T*_ (*α*_*F T*_ set by the fluctuations of *V* during the first 50 msec of the trace, before any stimulus), and if so defining a reference time *T*_*F T*_ as the latest time for which 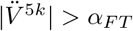 (Figure 5 Fourth row)):

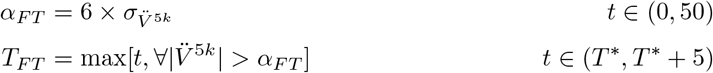

The start of the “pure” cell response 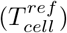 and the subsequent start time for the fitted data 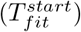 are then determined. If a fast transient was detected, then 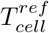 is defined by *T*_*F T*_ ; if not then by the transition time *T* ^*^ (the 1kHz low pass filtered voltage trace (Figure 5 Third row)). 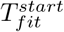 is then delayed relative to 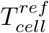 to minor the electrode component (assumed upper bound of 0.5 msec for the time constant), and to take into account any biphasic response from an over-compensated bridge, thus considering the delay 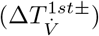 after 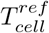 when 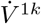 first becomes positive (negative) for a positive (negative) current transition.

Thus, 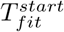 is given by 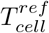 plus the maximum between 0.3 msec (ref. electrode time constant), and twice 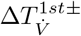 (ref. biphasic transition):

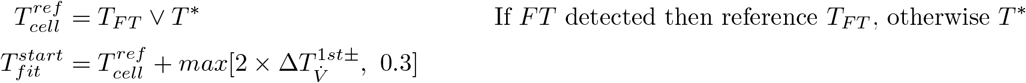

A single exponential plus constant fit *V*_*fit*_ is obtained for 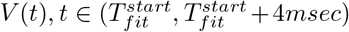, thus the initial 4 msec of the “pure” cell response following the bridge artifact (Figure 5 First row):

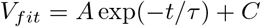

The estimated voltages immediately before and after the transition (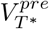 and 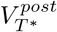), thus the relevant voltage step Δ*V* for the bridge error, are defined as follows:

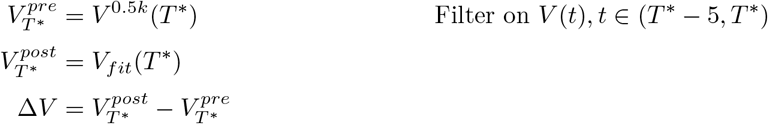

The bridge error *R*_*err*_ is given by:

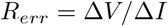

Finally, if any of the following conditions holds, then the estimate *R*_*err*_ is rejected:

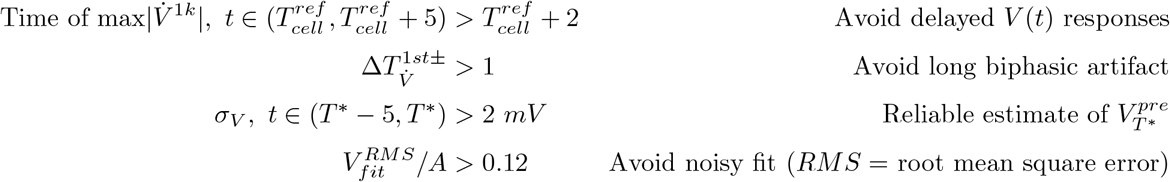

Within the time frame of a given set of trials as described previously, e.g. typically less than one minute, we assume that there is a correlation of bridge errors. As such, any rejected bridge estimate for trial indexed by *j* in a given set was replaced by a linear interpolation from flanking trials *i* and *k* with a valid *R*_*err*_ (*i < j < k*):

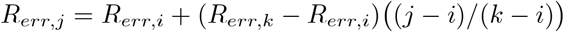

If there are no valid estimates flanking both sides of trial *j, R*_*err,j*_ was replaced by the mean of the accepted estimates over the whole set of trials.

To summarize, this *a posteriori* method for the estimation of the bridge balance, by being directly implemented in the analysis pipeline and performed on each recording allows to ensure that before performing any analysis on the current-clamp recordings traces, the voltage traces accurately reproduce the dynamic of the membrane upon the application of a stimulus. Indeed, we saw that uncompensated bridge balance induce a shift in the recorded membrane potential, which later could induce erroneous estimation of diverse neuronal properties like spike threshold, membrane resting potential, or input resistance. However, these different properties are key in various aspect of neurosciences, and therefore justifies the careful estimation of such experimental artifact. Furthermore, while bridge balance may be compensated directly during the recording session, and therefore in the published databases, like the Allen Cell Type Database, it may not be the case for other database which would require such a posteriori compensation. Furthermore, because of the additive nature of the bridge balance, it is coherent to systematically apply this procedure to any database in the pipeline, even to those already corrected, to ensure similar data treatment across the different databases before being able to compare and use the data together.

#### 3.3.2 Applying a common set of quality criteria

The TACO pipeline has been designed as a tool to study the electrophysiological properties of cortical neurons, as a function of the different neuron’s characteristics like the cortical areas, the layer, the cell type… As we saw before, this supposes the capacity of studying together electrophysiological recordings coming from different databases, and therefore the assurance that the data are equally treated before analysis. This aspect notably applies to the experimental artifacts potentially present in the databases, as we saw for the bridge balance. However, while the bridge balance is a common electrophysiological artifact justifying being specifically and systematically taken care of, the pipeline is meant to be used by other lab, also for future databases. Thus, the relevance of hard coding a set of quality criteria directly into the pipeline is questionable, as no consensus exists about what criteria must be applied to guarantee the data are of sufficient quality to be part of the analysis.

Therefore, to conciliate the subjective nature of quality criteria that would ensure viable analysis, the necessity to apply the same set of quality criteria to all databases so their data can be analyzed together, and the easiness of use of the pipeline, we designed it so that any lab can propose its own set of quality criteria, without any need to manually modify the pipeline to implement them into it. The goal of the pipeline notably being to characterize the firing properties and the Input/Output relationship, these quality criteria should allow specifying which of the sweeps present in the data can be considered for this firing analysis. Therefore, before applying them, the different aspects of the sweeps are characterized (see section Sweep analysis).

##### 3.3.2.1 Sweep analysis

###### 1 Defined protocol and trace id

As indicated in the part 3.2.2 Relying on limited number of information, the cell-sweep csv file contains for each cell the different sweeps to consider for the analysis. If multiple protocols have been performed, then the sweep id can be composed of both the protocol id and trace id (e.g.: Sweep id = [Protocol id] [Trace id]). Then to automatically detect protocol and traces id, the sweep id indicated is parsed by underscores (). The pre-last element is considered as the protocol id and the last element as the trace-id. If no underscore is detected in the sweep id, then the protocol id for every trace is marked as 1, and the trace-id corresponds to the sweep id.

###### 2 Holding current and applied stimulus current

The holding current corresponds to the current applied to the cell before and after the stimulus onset and offset times respectively, while the applied stimulus corresponds to the current applied in between. The stimulus amplitude therefore corresponds to the difference between the two. For each trace, the holding current and the applied stimulus were estimated by fitting a double Heaviside function to the filtered (2nd order Butterworth filter, LPF 1kHz) stimulus trace. To determine the weight of the fit, the stimulus derivative trace was computed. For each time point, the weight was computed as follow 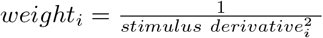 If more than half the weights values are infinity, then all weights are set to 1 (e.g.: in case of a synthesized current trace). Otherwise, the inf are replaced by the maximum weight value observed that is not inf. The weights are then normalized to the maximum weight value. The stimulus amplitude is then computed as the difference between the applied stimulus current and the holding current.

###### 3 - Bridge error analysis

The bridge balance analysis is performed as described above. However, if the analysis was not considered, notably for not respecting any of the late conditions, then the bridge balance for that sweep is linearly extrapolated inside a same protocol from the closest stimulus amplitude sweep for which a Bridge Error have been computed. To ensure consistent estimation across the traces and avoid any aberrant measure, extreme outliers of Bridge Error (if any) are removed (defined as values higher than Q3+3*IQR or lower than Q1-3*IQR, with IQR=Q3-Q1)

###### 4 - Sampling frequency

The trace sampling rate was computed from the time trace, by computing 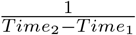

###### 5 - Sweep-based analysis of linear properties

For this analysis to be performed, several condition must be respected to ensure to be in the linear cell regime and avoid aberrant values. The conditions are as follow: 1) During the first half of the stimulus time window, no spike must be detected (following the procedure defined in the spike analysis paragraph). 2) The absolute value of the coefficient of variation of the voltage trace in the time window [Stimulus_onset_ − 200*ms*; Stimulus_onset_ − 5*ms*], as well as in the time window [half stimulus time window; Stimulus_offset_], must not exceed 0.01. 3) The absolute value of the stimulus amplitude must be higher than 2pA.

Once these different criteria have been assessed, the voltage trace is fitted to an exponential function as 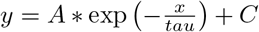 during a 300ms time window starting at the stimulus onset time. If the goodness of fit, defined as the RMSE of the fit normalized to the voltage amplitude (the voltage difference between the stimulus time window and before the stimulus) is higher than 0.3 then the input resistance, the membrane time constant, the resting potential, and the steady state potential are not defined for this sweep. Otherwise: the time constant is defined by the fit resulting value for tau. The holding potential is defined as the median value of the voltage trace before the stimulus onset. The steady state potential is defined as the offset (C) of the exponential fit. The input resistance is computed as follow:

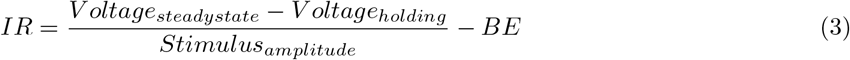

Finally, the resting membrane potential is given by :

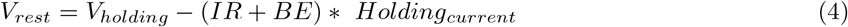

To ensure consistent estimation across the traces and avoid any aberrant measure, extreme outliers (if any) of time constant and input resistance are removed (defined as values higher than Q3+3*IQR or lower than Q1-3*IQR, with IQR=Q3-Q1).

The cellular input resistance is defined as the slope of a linear regression of the subthreshold potential steady-states and stimulus current amplitude. The cell’s resting potential is defined as the intercept of the linear regression of holding membrane potential and holding current. The cellular time constant is defined as the average of the sweep computed time constant.

The cautious sweep-related analysis is an essential step for the characterization of neuronal electrophysiology. All the previously defined sweep-related measurements, together with the sweeps raw potential and current traces can then be use to assess the viability of the different sweeps, and decide which one should be part of the firing analysis. However, as we explained before, there are no universal definition about what makes a trace suitable for analysis. Therefore, the quality analysis should be easily customizable.

##### 3.3.2.2 Automatic integration of custom trace based quality analysis

The key design principle of the TACO pipeline was ease of use, which is not immediately compatible with the exterior proposition of an unpredictable and variable set of quality assessment potentially requiring data pre-analyzed inside the pipeline. Therefore, we designed a solution so that any lab can propose a set of quality analysis that are to be performed sweep by sweep, without any need to manually encode these into the pipeline.

This process relies on the participation of external labs which provide a Python script describing the different analysis in the form of different functions to be performed sequentially. Simply by providing the name of the user’s script at the beginning of the analysis, the TACO pipeline will integrate the different functions, and apply each of them on each cell’s sweep. However, as stated before, these different analysis are to be performed sweep by sweep, and so based on experimental values specific to the sweep at stake (resting potential, bridge balance, recorded traces,…). Therefore, to ensure the functions receive the appropriate values, and that the quality analysis is tractable, the TACO pipeline sequentially makes the different values computed in the last part, as well as the related potential and current traces, available as global variables in a side module, that is to be imported in the user-written script. Thus, to access these different values, and perform any kind of analysis, the functions simply need to refer to them, as if they were already defined in the script. Finally, the results of each of these quality assessments will be stored as part of the global cell analysis, enabling a tractability of the quality analysis.

By proposing a way for any user to define a set of quality analysis that will be applied on each cell’s sweeps as if these user-defined function were natively integrated into the pipeline, the TACO pipeline reconciles the need to perform common quality analysis on all the data before being able to reliably compare different database, with the easiness of use of the pipeline, by solely requiring to write functions as if they already were in the pipeline. Yet this approach still presents some drawbacks, that would need further development to be erased. The first of these is that while the functions written by the user can directly call the required TACO-computed variables, this step requires that the variable are written as they are in the TACO pipeline, making it case-sensitive. To overcome this, we provide a template script containing a description of the different variables available for the functions, therefore providing a reference, but not guarantying it from misspelling, and therefore skip of the related function. The second drawback is the fact that the different quality analysis are used to determine if a sweep should be part of the firing analysis, but in an all-or-none fashion. In other word, the failing of any of the user defined function would be sufficient to reject the current sweep. Yet, we can easily think of conditions that would need to occur simultaneously to be considered as prohibitive for the sweep to be considered. For now, such configuration should be part of the same function and render a single quality assessment. Finally, for the moment the spike-related features are not available yet for quality analysis, while they could be used to estimate if a sweep should be part of the firing analysis. As explained before, this quality analysis aimed at providing a convenient way for external user to propose a set of quality tests, as it is crucial to have a common quality analysis for all traces of any databases, before being able to characterize their electrophysiological properties, and notably the dependence of the firing response of the neuron regarding the input it receives.

### 3.4 Detailed analysis of firing properties

The electrophysiological characterization of a neuron comprises a range of aspects, from subthreshold dynamics to spike generation and transfer function description. These different aspects can be assessed from a classical current-clamp experiment, by applying multiple levels of current stimulation, and observing the neuron’s response. While different important aspect of neuronal physiology can be assessed during the sweep analysis, like the input resistance, the resting membrane potential or the time constant (see section Sweep-based analysis of linear properties), the firing properties compose the super-threshold response of the neuron. To be able to appropriately characterize the input – output relationship of a neuron, it is necessary to first be able to identify spikes in recording traces.

#### 3.4.1 Spike analysis

As stated before, action potential generation constitutes one main aspect of neuronal response to external stimuli. Characterized by an abrupt depolarization immediately followed by a hyperpolarization of the neuron’s membrane, spikes detection is a crucial step toward characterizing neuronal population dynamics. While most of commercial recording softwares propose built-in spike detection algorithms, numerous open-source solutions also propose custom methods for spike detection (see section Diversity of electrophysiological data acquisition and analysis environment). Multiple solutions have been designed, notably relying on clustering analysis to separate the spike waveform from the other neuron’s signal components (Balasubramanian and Obeid, 2011; Toosi et al., 2021; Pachitariu et al., 2024) or spike feature distribution (Bestel et al., 2012). Such complex solutions are needed especially for extracellular recording experiment, with electrodes recording Local Filed Potential (LFP) signals originating from multiple neurons. In the context of the TACO pipeline, databases of interest are composed of single neuron, intracellular recordings, making it simpler to detect single action potential. Therefore, we used a traced-based method for the detection of spikes in voltage traces, as well as the characterization of diverse spike-related features, as described in Gouwens et al. (2019).

The spike detection is made as follow:

Putative spike threshold are detected by detecting in the voltage trace times where 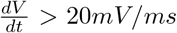. The putative spikes’ peak are then detected as the maximum potential values between two consecutive putative spike thresholds. To ensure this method only detects actual action potentials, and not fast noise-induced membrane depolarization, each putative spike was validated by asserting that the potential difference between the peak and the threshold was at least of 20mV, and that the peak potential was higher than 30mV. Any putative spike for which one of these two conditions was not observed, was not considered as a spike for the rest of the analysis.

Spike upstroke (and downstroke) were defined as the maximum (minimum) value of the first potential trace time-derivative between the threshold and the peak (between the peak and the next threshold. The spike thresholds were then redefined as the last time before the spike upstroke for which 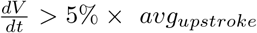, with *avg*_upstroke_ corresponding to the average value of the potential first time-derivative at the time of the upstroke. Then, the alternation of spikes’ thresholds and peaks were checked to ensure no overlapping (i.e.: ensure no consecutive thresholds or peaks). In this case, the two events were merged.

The spikes’ trough were then defined as the minimum potential values between a spike’s peak and the following spike’s threshold. A fast after-spike hyperpolarization (fAHP) was defined for a given spike, if during the 5ms time window following a spike’s peak the time derivative of the membrane potential went positive, and the membrane potential went lower than the spike’s threshold. An after-spike depolarization (ADP) was detected by detecting between a spike’s downstroke time and the next spike threshold the first occurrence where 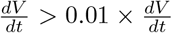 at spike downstroke (*t*_*A*_). Then between this time and the next spike threshold, the first occurrence where 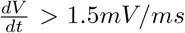 was detected (*t*_*B*_) and use as a lower time limit to detect the first time where the potential first time derivative went negative. For the potential ADP to be validated, the following criteria must be observed: 1) the time difference between *t*_*A*_ and *t*_*B*_ must be lower than 2ms, 2) the potential difference between the ADP and the minimum potential between the ADP and next spike threshold must be higher than 1mV, 3) the potential difference between ADP and the potential at *t*_*A*_ must be lower than 10mV, and 4) the time difference between ADP and *t*_*A*_ must be lower than 5ms.

This traced-based spike analysis allowed the systematic detection of spike as well as the precise timing of each spike feature. Therefore, additionally to peak, threshold, upstroke, downstroke, fAHP, trough and ADP, for any given spike, spike height is defined as the difference of potential between the peak and the threshold, and spike width at half height, is defined by the time difference between *t*_*h,d*_ and *t*_*h,a*_ with *t*_*h*_ the time when the membrane potential reaches half the amplitude (defined as *v*_*amp/*2_ = *v*_*th*_ + 0.5 · *Spike*_*amp*_) in the ascending part of the spike (i.e., *t*_*h,a*_) and in the descending part of the spike (i.e., *t*_*h,d*_). The detection and characterization of spike-related features represent crucial steps for characterizing the cell’s response to a stimulus, as well as the neuron’s input-output relationship.

#### 3.4.2 Consistent quantification of I/O relationship

As we explained before, the characterization of I/O relationship can highly vary between labs, due either to difference in experimental procedure, or to the fitting methods used, which can lead to highly different quantification of I/O properties for similar data (Figure 1). For these reasons, a single common method to describe continuous-like neuronal transfer function could enable better interoperability of electrophysiological data, and cross-studies comparisons of I/O features. As part of the TACO pipeline, we implemented an I/O fitting procedure robust to the difference in methodological procedure employed during the recording experiment and able to characterize the diversity of I/O relationship shape. We also defined methods to obtain reasonable estimates for initial conditions, so this method could be implemented in other frameworks.

#### 3.4.3 D IO Fit (Hill-Sigmoid)

To address response diversity we fit the data to a function *HS* composed of an ascending Hill type characteristic *H* multiplied by a descending logistic sigmoid *S*. Notably, the Hill-Sigmoid description reasonably captures both concave and abrupt initial portions of the I/O relation, different curvatures between the initial portion and any subsequent convex portion, and any eventual decay or failure of the response for large inputs. In the one-dimensional case, the independent variable *x* is either the current stimulus *I* or the relative excitatory conductance stimulus 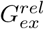:

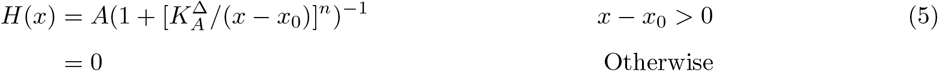

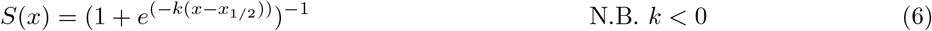

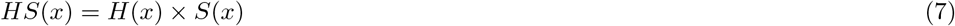

Given the many free parameters of the target function, the fit procedure relied crucially on their initial conditions and constraints on their minima and maxima. These parameters were derived from an initial polynomial fit to the data that captures the overall envelope of likely I/O relations. We first consider the one-dimensional case:

1. Consider a set of *N* input/response pairs *R* = [(*x*_*n*_, *r*_*n*_)], ordered according to the input values, defining the stimulus and response extremes (*x*_*min*_, *x*_*max*_, and *r*_*min*_, *r*_*max*_) over the pairs. The essential dynamic portion of the stimulus-response data, *R*_*θ*_, is obtained from *R* to exclude both small or zero responses, corresponding to essentially sub-threshold input, on the left, and a possible falling (or failure) response for large inputs on the right:
  - For *n* = (0, *N* − 1), let *n*_*ref*_ correspond to the first time *r*_*n*_ ≥ (*r*_*max*_*/*10). Set 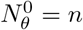 for the last time *r*_*n*_ = 0 (*n < n*_*ref*_); if no zero response in this part of the data, then set 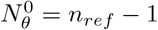.
  - For *n* = (*N* − 1, 0), set 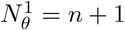 the first time *r*_*n*_ ≥ (*r*_*max*_*/*2) The indices 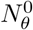 and 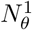 anchor the trimmed data to subthreshold responses, and set the domain of *R*_*θ*_ as follows:

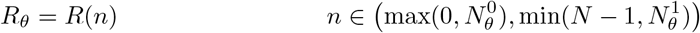 The limits on the beginning and ending values for *n* account for values of 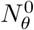 or 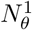 at the limits of *R*.
2. Two properties of *R*_*θ*_ are then verified before fitting can continue. First, to avoid overfitting, there must be at least 4 non-zero *r*_*n*_ values in *R*_*θ*_. Second, to avoid sparse sampling of the input space, a sufficiently spread out distribution for fitting is validated by dividing the domain of *R*_*θ*_ into 8 identical segments, and requiring that at least 4 include a stimulus-response pair. If these conditions are met, a third-order polynomial *P* is then fit to the trimmed data *R*_*θ*_. If the peak of *P* occurs before the end of *R*_*θ*_, thus including a descending component, then the stimulus-response pairs are fitted to the composite *HS* function; otherwise the fit is to the Hill function *H* alone.
3. We first derive the fitting parameters for *H* from *P*. Defining *P*_*min*_ and *P*_*max*_ as the minimum and maximum values of *P* over the domain of *R* (*x*_*min*_ ≤ *x* ≤ *x*_*max*_), respectively, the initial values and constraints for *A* are:

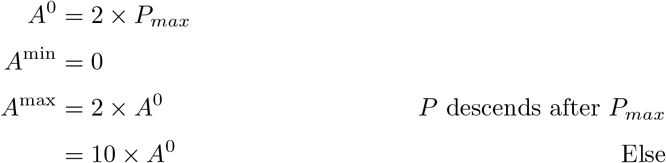 The first upper bound on *A* avoids pathological overshoot of the fit to sparse data; the last upper bound on *A* avoids overfitting in the event of an essentially linear relation between stimulus and response.
Let *P*_up_ be the ascending 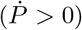 segment over the domain of *R* that immediately preceeds *P*_*max*_, e.g. ignoring any initial descending portion of *P*. If the form of *P* suggests a threshold input within *R*, specifically if either *P*_up_ intersects the X axis within the domain of *R*, or (more implictly) if *P* descends at the start of 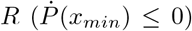, then the initial value 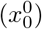 for *x*^0^ is given by the first entry of the trimmed data:

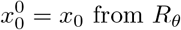 Otherwise (thus no subthreshold component), an estimate for *x*^0^ is provided by a linear extrapolation backwards from *P*, evaluated at the start of *R*:

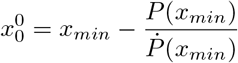 The constraints on the fit values of *x*^0^ are:

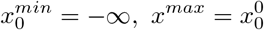 The fitting parameters for *n* are:

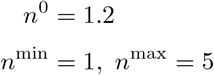 The initial guess of 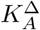 corresponds to the distance between 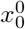, and the input corresponding to the presumed mid-point of the output:

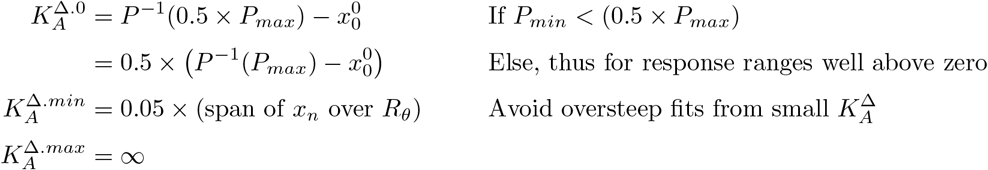 Note that the second definition of 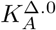 takes into account a data set *R* without a subthreshold component; indeed some definitions of the response have a non-zero minimum response *a priori*. Finally, 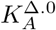 is then bounded by 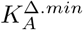 as necessary.
4. If indicated by a descending phase in the data, the fitting parameters of *S* are derived from *P* after *P*_*max*_. A linear fit (*L*(*x*) = *ax* +*b*)) is obtained for *P* between *P* ^−1^(*P*_*max*_) and the final input in *R*_*θ*_. The initial value for the sigmoid midpoint *x*_1*/*2_ is set to where *L*(*x*) descends to half it’s value at *P* ^−1^(*P*_*max*_):

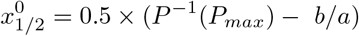 There are no constraints on *x*_1*/*2_. The fit parameters for the sigmoid steepness *k* are given by the slope of *L* normalized by it’s value at *P* ^−1^(*P*_*max*_):

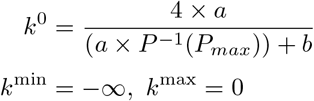

#### 3.4.4 D IO Measures: Threshold, Gain, Saturation and Failure

Consider an input output characteristic ℱ resulting from the fit to either *HS* or *H*. Let ℱ_*max*_ be the maximum value of that fit over its entire domain, and ℱ^*R*^ correspond to ℱ over *R*.

The threshold 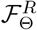 is defined as the first input value in ℱ^*R*^ corresponding to the minimum detectable measure of the response variable, e.g. for the response given by the spike frequency over 500 milliseconds, this corresponds to a single spike, thus 2Hz.

The gain 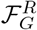 is estimated at the middle of the assumed dynamic range of ℱ^*R*^ given by its minimum and maximum values (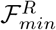and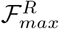). 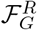 is defined as the slope of a linear fit of the rising phase of ℱ^*R*^, *x* ∈ (*x*_*start*_, *x*_*end*_), over the middle 25% of its range:

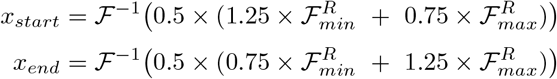

Saturation 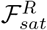 is only defined when either the ratio of the final slope of ℱ^*R*^ and ℱ_*G*_ is ≤ 0.2, or 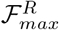 is ≥ 0.75 ×ℱ_*max*_; in either case 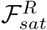 is then set to ℱ_*max*_. Note that the latter condition holds when ℱ^*R*^ includes a descending portion.

Response failure is detected when 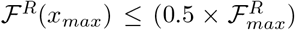, and if so then the failure input 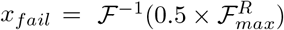 within the descending phase of ℱ^*R*^.

The careful data-based estimation of the different parameters’ initial conditions and boundaries, allowed for a single procedure able to carefully fit a wide diversity of Type I-like I/O relationship to either a Hill (H) or a Hill-Sigmoid (HS) function. Then the precisely defined method for extracting I/O features from it makes this method a powerful way of characterizing neuronal transfer function from current-clamp experiments (see Figure 1 Hill-Sigmoid fit). Such reliable and detailed method is crucial for the interoperability of electrophysiological database and for the study of neuronal firing properties

#### 3.4.5 A global method to quantify adaptation

In the context of current-clamp experiments, in addition to the diversity of firing behavior observable inside a population, the sampling of input space can greatly vary between databases, making the cross-database comparison of SFA complicated. To overcome the database-related variability due to non-standardized recording procedure (i.e.: different number of recorded traces, different input amplitude, different response duration…), we designed a method to quantify the adaptation of different spike-related features (e.g.: spike amplitude, ISI, spike width at half height…) based on measures from all available traces. We fit the evolution of the measure of interest to an exponential plus constant function, using spike index as the independent variable to make the analysis more robust. Indeed, as spike index monotonically increases with spike time, the time-dependent adaptation mechanism is considered in the analysis, while reducing the effect of precise spike timing. Furthermore, as the precise quantification of the spike property may be impacted by the amplitude of the input current applied, we normalize the measures of interest in a specific trace by the value at the first spike, thus considering adaptation as a relative mechanism, rather than absolute. Our quantification of the adaptation ultimately consists in the comparison of a constant and modulated response component (Figure 7).

**Figure 6:**
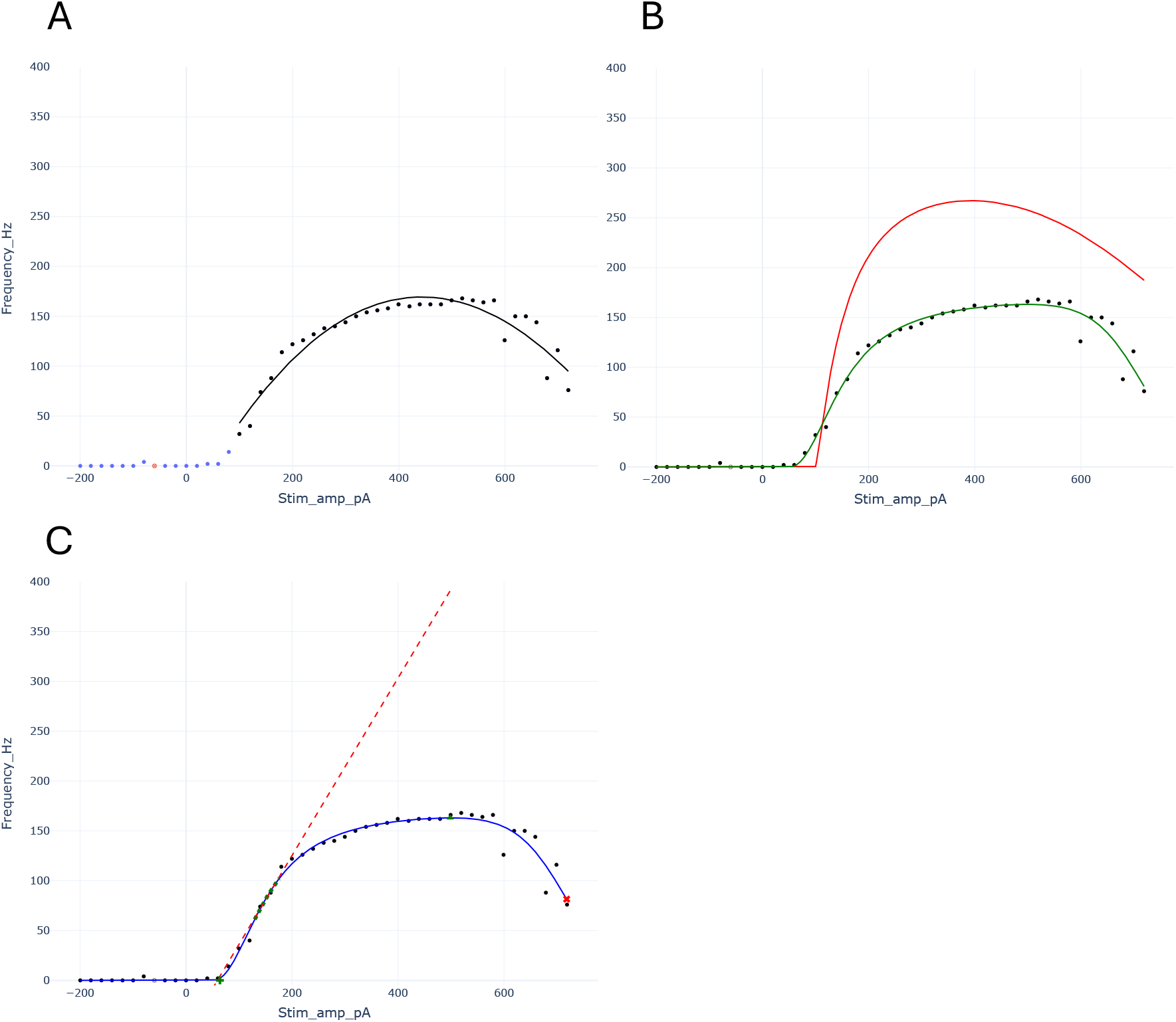
Detailed example of I/O fitting procedure to Hill-Sigmoid function. **A**: Fit of third order polynomial (black line) to trimmed stimulus-frequency data (black dot). Blue dots represents sweeps below the trimming threshold, and are not considered for this step. Red dot represents a sweep which failed QC analysis, therefore not considered for the firing analysis. This fits catches the global shape of the I/O relationship and notably the descending portion. **B**: Fit of a Hill-Sigmoid function to full stimulus-frequency data. The red curve represents the Hill-Sigmoid curve with initial fit conditions, and the green the Hill-Sigmoid curve after the fit to the stimulus-frequency data. **C**: Measurement of I/O properties from the Hill-Sigmoid curve (blue). The gain is obtained by performing a linear fit (red dashed line) to the middle portion of the ascending phase of the Hill-Sigmoid curve (green dashed line). The threshold is obtained as the minimum stimulus value corresponding to the minimum observable response on the Hill)-Sigmoid curve (green cross). The Saturation is represented by the maximum value of the Hill-Sigmoid curve (green triangle) and the response failure (which is identified if the Hill-Sigmoid curve become inferior to half the saturation frequency after the saturation is represented by the red cross. This example is taken from sweep CC_5_12, cell 2017_10_05_2_6 from Lantyer Database.

**Figure 7:**
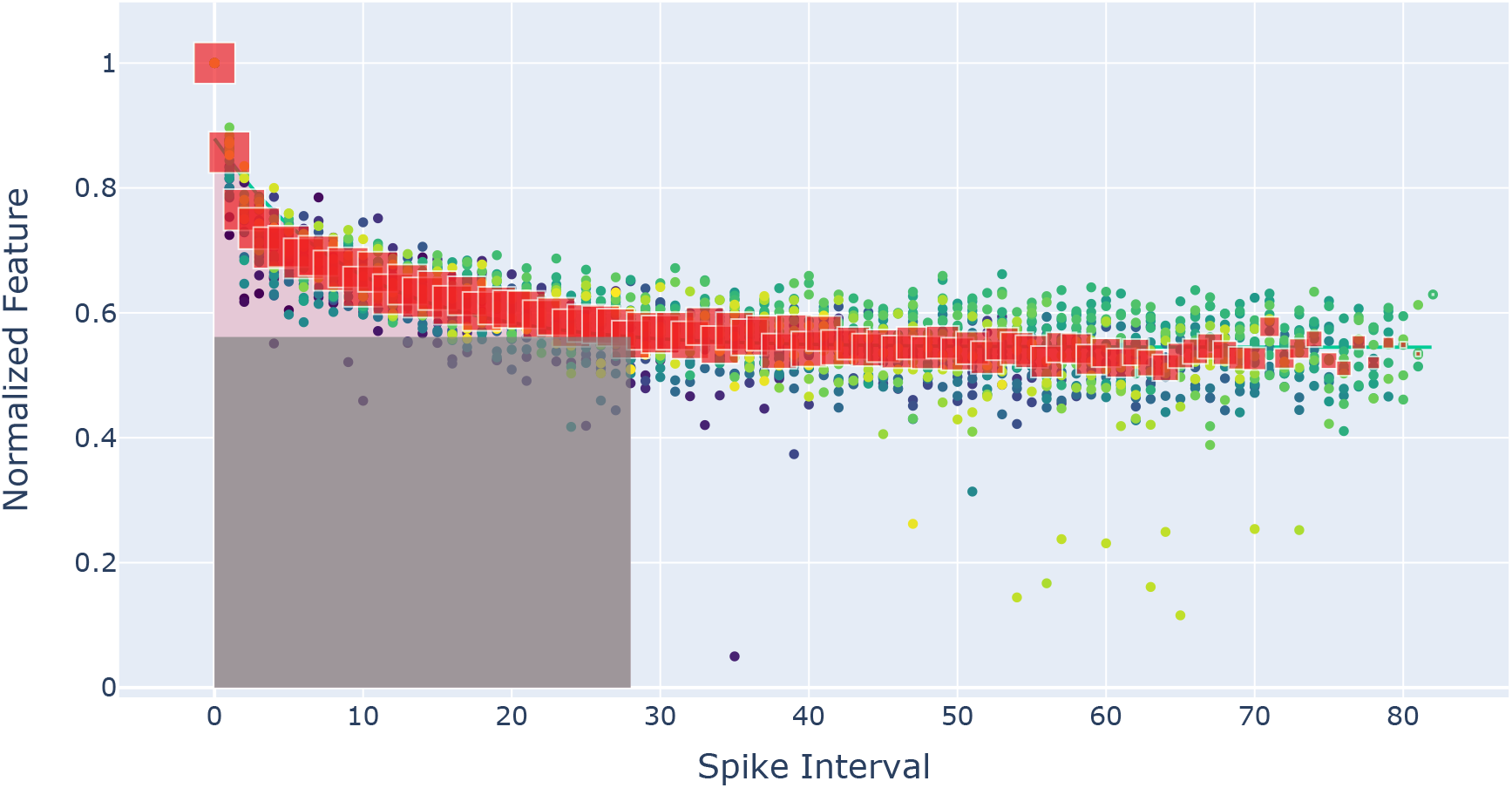
Example of Instantaneous frequency adaptation computation. For each sweep, the instantaneous frequency for each spike index is computed (individual point, the color represent the stimulus amplitude from blue 80 pA to yellow 718 pA). The red squares represent the median value for each spike index, the size represents the number of value across sweeps for each spike index (from 33 observations at spike index 0 to 1 observation for spike index 82). The median values are used to fit an exponential curve (green line). The grey area represents the constant component of the response, when the instantaneous frequency does not vary anymore. The pink area represents the modulation component. The adaptation index is computed as the relative proportion of modulated component compared to the addition of both the constant and modulated components of the response. This example is taken from sweep CC_5_12, cell 2017_10_05_2_6 from Lantyer Database.

Precisely, we consider a function *F*_*n*_ considering the measure of interest at spike *n* (if the measure of interest considers the spike interval, like for instantaneous frequency, it is defined by spike *n* and *n* + 1). For each response trace *i*, we collected the *F*_*n*_ values over the response duration, adjusted to the first spike value (for 0-bounded values like frequencies, 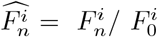, for non-bounded measure like voltage-measure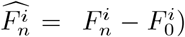. To reduce the impact of outliers, an exponential plus constant 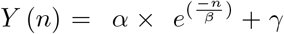 is fitted to the median of the 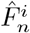 values over all traces, with regard to the spike index *n*, weighted by the number of observations *ω*_*n*_ at spike n over all traces. The only constraint is for the adaptation constant *β* which is high-bounded by *N*_max_ (maximum number of spike over all traces). If 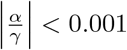, then the response is considered as constant given by the average value of the fit, thus: *Y* (*n*) = *Y* (*n*). Then the positive part of the fit is retained as *Y*_+_ (*n*) = max[0, *Y* (*n*)] to avoid considering possible negative values.

We then define *N*_*A*_ as the maximum spike index for which we could evaluate the adaptation, such that *N*_*A*_ = min[30, *N*_*max*_]. The constant component of the response C is defined by *C*_*ref*_ which is the minimum value of *Y*_+(*n*)_ over the window. The area of the constant component is then computed as *C* = (*N*_*A*_ − 1)* *C*_*ref*_. For the modulated part *Y* ^*M*^ (*n*) of the fit, it is obtained by subtracting its final value in the window such as : *Y* ^*M*^ (*n*) = *Y*_+_ (*n*) − *Y*_+_ (*N*_*A*_ − 1). The area of the modulation component M, is given by the absolute value of the of the area of *Y* ^*M*^ (*n*), such as 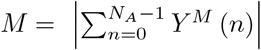.

The adaptation index is thus defined as 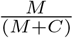, allowing to describe the relative variation of a given measure with regards to a constant part, for both increasing of decreasing values (e.g.: spike width or instantaneous frequency).

### 3.5 Using the TACO pipeline

All of the required python scripts are available for download at : https://github.com/Jballbe/TACO_pipeline. To ensure convenient use of the TACO pipeline, the different steps involved in running and analyzing the analysis have been implemented in a Python-based graphical user interface (GUI), utilizing the Shiny suite for Python. Once the different python packages and TACO pipeline scripts are downloaded, the user can launch the application from the command line, using the following command after entering the directory in which the scripts are downloaded:

~~~
shiny run --reload --launch-browser TACO_pipeline_App.py
~~~

The application then automatically opens a tab in the internet browser.

#### 3.5.1 1 - Create the JSON configuration file

To ensure the proper functioning of the TACO pipeline, several pieces of information need to be provided, particularly to specify the locations of user-defined sweep QC scripts and trace-extraction scripts. Additionally, the Cell-Sweep and Population Class tables required for each database need to be specified. (See section 3.2.2 Relying on limited number of information.)

To validate the entered paths, a check can be performed to ensure they exist. A visual feedback mechanism allows users to monitor the construction of the JSON configuration file. If an error is detected in any information for a database, users can re-enter the information for that database, and the JSON file will be corrected.

Once all the information is entered, users can press the “Save JSON” button to save the file to the desired location.

#### 3.5.2 2 - Run analysis

Once the JSON file has been created, the user can run the analysis which relies on the previously defined JSON configuration file. Before launching the analysis, the user can select the step of the analysis to perform (e.g.: Sweep analysis, Spike analysis, Metadata, Firing analysis or all at once), and choose wether to overwrite already exciting analysis files. The analysis can then be performed using parallel processing. In this case the user can enter the number of processing to allocate for this analysis. Once launched, the user can follow the progress of the analysis database per database.

#### 3.5.3 3 - Cell visualization

Once the analysis is done, the user can investigate the results cell by cell. For each step of the analysis described above (i.e.: spike detection, bridge error estimation, linear properties quantification, fit of the I/O relationship and spike frequency adaptation), an interactive plot presents the data processing and resulting quantification. In addition, different tables gather sweep-related information (i.e.: stimulus amplitude, resting membrane potential…), as well as results from the quality criteria step. Implementing a cell viewer application was necessary to enable visual inspection of the different part of the analysis and keep a critical perspective on the different solution we came up with (see Supplementary figures 13, 14, 15, 16, 17).

## 4 Result for diverse databases of electrophysiological recordings

### 4.1 Presentation of the gathered databases

#### 4.1.1 Allen Cell Type Database (Gouwens et al., 2019)

The Allen Institute has proposed for the last couple of years various resources to support open science in the neuroscientific field. Indeed, by publishing multiple open access databases of diverse neurological data, from transcriptomal studies, connectivity maps, anatomical atlas to electrophysiological characterization and neuronal modeling, in human, non-human primates and mouse, it represents a key resource in the neurosciences field, supporting diverse types of research (Ghaderi et al., 2018; Teeter et al., 2018; Billeh et al., 2020; Gouwens et al., 2020; Yao et al., 2021). The Allen Cell Type Database presents a detailed database of electrophysiological recordings, targeting all cortical layers, using different transgenic mouse line, mainly from visual cortex but it contains some cells originating from other cortical areas like auditory, retro-splenial, or somatosensory area. Each cell presented in this database is presented with different anatomical details like its location (cortical area, sub-area, cortical layer), and physiological information (i.e.: transgenic mice lines, was the cell expressing the reporter of interest, …). This different information represents key neuronal characteristics and are notably of importance when studying specific cell type classification (Ghaderi et al., 2018; Rodríguez-Collado and Rueda, 2021). Furthermore, the detailed electrophysiological characterization, using diverse stimulus conditions enabled the definition of diverse electrophysiological, morphological and electro-morphological classes (Gouwens et al., 2019), as well as the generation of neuronal models embracing electro, morpho and transcriptomic cell characteristics (Nandi et al., 2022). To fully embrace open science requirements, this database has been released in NWB file format, and a detailed analysis framework and online support helps the user to deeply analyze the data.

#### 4.1.2 Lantyer Database (da Silva Lantyer et al., 2018)

Presented in 2018, this database of whole-cell intracellular recordings specifically targeted cells from layer 2/3 in the somatosensory cortex. Its reuse potential notably resides in the diversity of the stimulus applied on the cells, with both voltage-clamp and current-clamp recordings, with different type of inputs applied like “step and hold” (for current-clamp), “Frozen-noise” (for both current-clamp and voltage-clamp) or Sawtooth (for voltage-clamp). Published on GigaScience repository using Matlab file format, the current-clamp data acquired using step-and-hold protocol are appropriate candidate for the TACO pipeline, especially as the data presented seems not to be bridge compensated.

#### 4.1.3 Harrison Database (Harrison et al., 2015)

In 2015, Harrison et al., 2015 introduced a database of patch-clamp recordings of pyramidal cells originating from layer 2/3, 4 and 5 in somatosensory cortex. Using either “square-pulse” or “naturalistic fluctuating stimuli”, they characterized different subthreshold, firing and action-potential-related properties according to the cell class (Layer 2/3, Layer 4, thick-tufted or slender-tufted Layer 5). They implemented a method to generate from these different measurements, single-cell fitted refractory Exponential Integrate-and-Fire (rEIF) models, with the aim of implementing network models of heterogeneous population. The data base has been Published on Datadryad repository, using binary file format.

#### 4.1.4 Scala 2019 database (Scala et al., 2019)

To address the connectivity as well as the cellular morphological and electrophysiological differences between cortical areas, Scala et al., 2019 performed patch-seq recordings of cells originating from layer 4 and layer 5 in either visual or somatosensory cortex, using difference transgenic mice lines. While some cells were directly genetically targeted, other where morphologically targeted and their cellular type were inferred from their morphology. More precisely, large, small, double-bouquet and horizontally basket cells were considered as paravalbumin (Pvalb) expressing cells, bipolar cells with small soma extending to L1 and L5 were considered as vasopeptide (Vip) expressing cells, while non-Martinotti cells from SS cortex were considered as somatostatin (Sst) expressing cells (while in the visual cortex Sst interneurons are reported be predominantly Martinotti cells). It was published on the Zenodo repository using Matlab file format.

#### 4.1.5 Scala 2021 database (Scala et al., 2021)

As part of the BRAIN initiative cell consensus network Scala et al., 2021 presented a database to characterize the neuronal transcriptomic diversity in motor cortex. They recorded under patch-seq technique cells from various transgenic mouse line, in different cortical layers at different recording temperature (room temperature, and physiological temperature). This *in vitro* study notably shown that morpho-electrical variability and transcriptomic variability were correlated, suggesting that transcriptomic types constitute a continuous landscape, rather than discrete t-types families.

#### 4.1.6 NVC database

The host laboratory constituted over the last years a database of in vivo whole-cell recordings originating from different animals (cat, mouse and rat). While non-targeted, this database presents the unique advantage of having cells recorded in both current and conductance clamp configuration. Such database represents a unique opportunity for establishing a correspondence between these two recording configurations, and thus infer conductance-based properties from current clamp recordings.

Using the TACO pipeline, we gathered multiple intracellular recording databases presenting a variety in both the technics employed and the specific cell targeted in the experiment. More specifically, we gathered six different databases, covering the main excitatory and inhibitory cell types (i.e.: Pvalb, Sst, Vip, Htr3a) present in the cerebral cortex (Xu et al., 2010; Rudy et al., 2011; Tremblay et al., 2016; Rodarie et al., 2022),in three main cortical areas (i.e.: Visual cortex, Motor cortex, and Somatosensory cortex), and covering across all cortical layers (Figure 8). Apart from these physiological differences, these databases notably differ from each other by there experimental paradigms, some being recorded *in vivo* (e.g.: NVC database) and other *in vitro* (e.g.: Allen cell type, Lantyer, Scala 2019 and Scala 2021 databases), or differing by the recording temperature applied during the experiment. Finally, the curation applied during the recording may greatly vary between databases, making the comparison between databases tedious (Figure 9).

**Figure 8:**
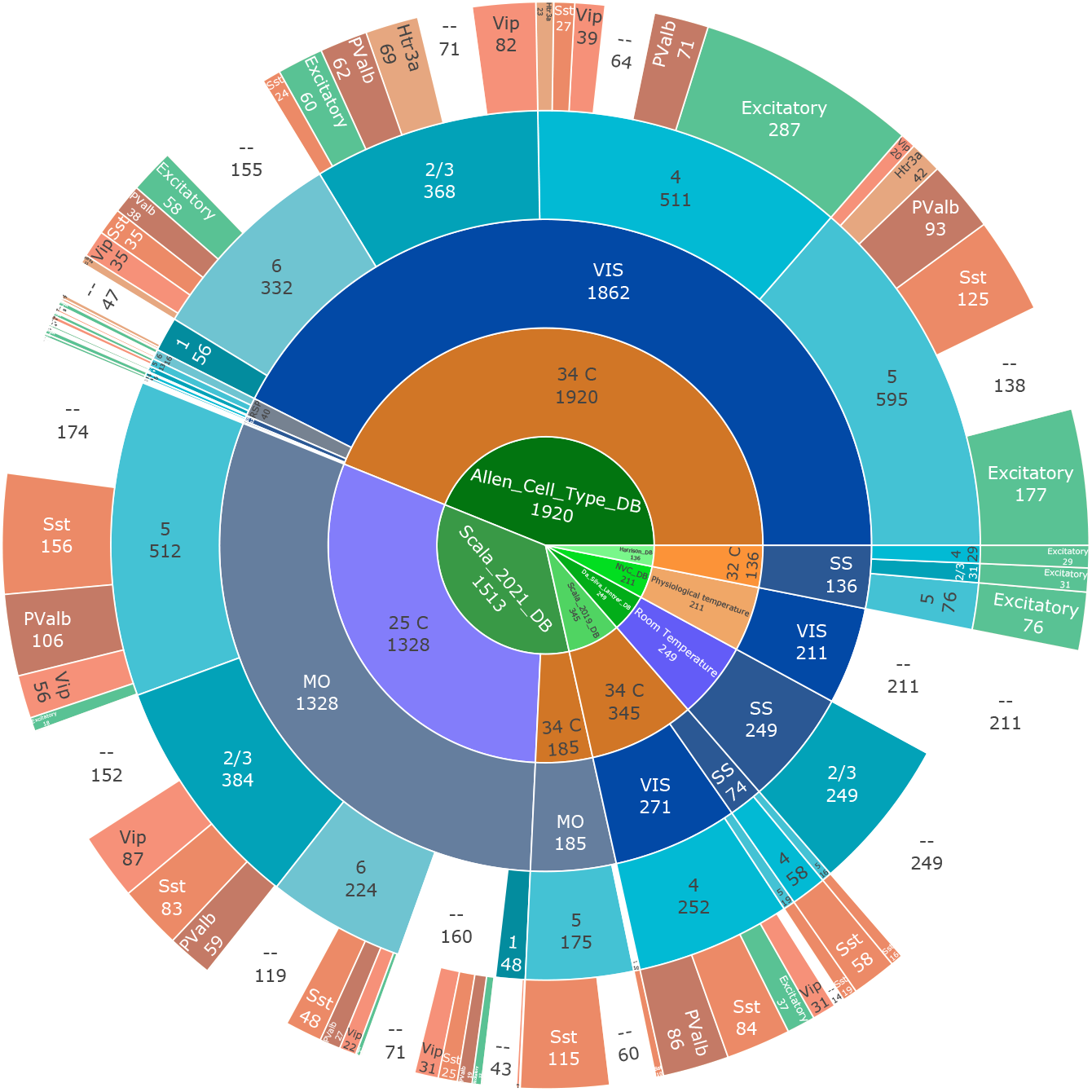
Main categorical repartition of databases. At the time of writing, we gathered six different databases of intracellular recordings. We gathered data recorded mainly in Visual, motor and somato-sensory cortex, in different recording temperature, covering all cortical layers. Furthermore, different databases made targeted recording allowing to know the cell type recorded (If a given information was not presented in the database, it was marked ‘–’)

**Figure 9:**
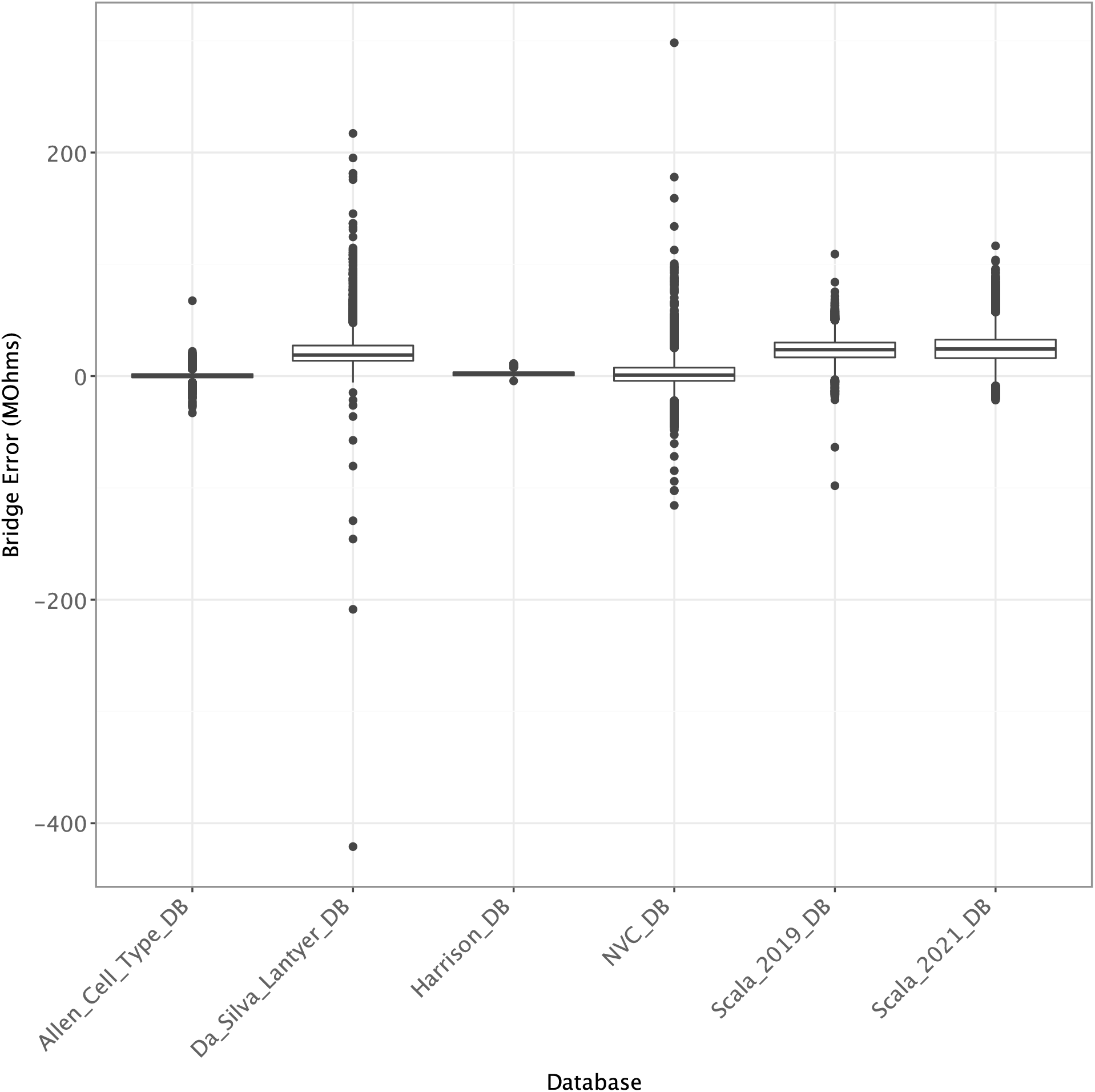
Distribution of *a posteriori* estimated bridge balance per trace in different publicly available databases. We measured the Bridge Balance *a posteriori* in different publicly available databases of intracellular recordings. These databases were made of excitatory cells recorded in rat somatosensory cortex (Harrison et al., 2015), inhibitory cells recorded in mouse somatosensory cortex (da Silva Lantyer et al., 2018), both excitatory and inhibitory cells recorded under various conditions in mouse visual and motor cortex (Scala et al., 2019, 2021), and the database provided by the Allen Institute (Gouwens et al., 2019). Finally, we also used a database from the host laboratory (NVC) composed of *in vivo* recording in mouse, rat and cat visual cortex.

### 4.2 Predictive performance of features extracted with TACO

A potential benefit of gathering different databases through the same pipeline, would be to rely on the electrophysiological properties computed similarly to infer other cellular properties, such as molecular identity, or their excitatory or inhibitory profiles. Indeed, several studied have tried to identify neuronal classes based on different electrophysiological properties. Notably, to decipher between PValb, Sst, and pyramidal (glutamatergic) neurons, (Ghaderi et al., 2018) relied on the discrete cosine transform of spontaneous *in vivo* spike waveform, and achieved average classification accuracy of 92.67±0.54% (Pvalb: 91.59±1.69%, Sst: 89.06±1.99, Pyramidal: 97.47±0.67%). They repeated the same procedure for *in vitro* cells, relying on data from the Allen Cell Type Database to extract numerous spike waveforms from 50 cells per type (PValb, Sst, Vip, Htr3a, and Pyramidal). Relying on a semi-supervised clustering algorithm, they achieved an overall accuracy of 84.13±0.81% (Pvalb: 93.57±0.59%, Sst: 89.15±0.63, Vip: 81.69±0.56%, Htr3a: 79.23±1.38%, Pyramidal: 77.02±0.91%). Similarly, Rodríguez-Collado and Rueda (2021) relied on the parameters of Frequency Modulated Möbius model derived from spike waveforms to establish a cellular taxonomy of Cre-lines, and decipher between common cell types (Pvalb, Sst, Htr3a, Vip, and Glutamatergic). They achieved an overall accuracy of 80.3% using model averaged neural network classifier, the best predicted class being glutamatergic cells (95.9%, PValb: 86.9%, Sst: 72.5%, Htr3a:54.9% and Vip:46.5%). More recently Ophir et al. (2024), additionally to providing a domain adaptation framework to co-jointly study mouse and human electrophysiological data, proposed the use of Locally Sparsed Artificial Neuronal Network (LSPIN) to identify classical neuronal subtypes (Pvalb, Sst, Htr3a, Vip, and Glutamatergic). Relying on a set of 41 electrophysiological descriptor per cell derived from the Allen Cell Type Database (Gouwens et al., 2019), they achived a global accuracy of 91.6% (Glutamatergic: 97.8% :Htr3a: 82.9%, Pvalb 86.8%, Sst 79.2% and Vip 95.8%), together with a mean F1-score of 87.7%. Using a more classical approach like a Random forest, the overall accuracy was of 82.4% and the F1-score of 72.6%.

#### 4.2.1 Within a database

To test the predictive potential of the feature extracted using the TACO pipeline, we trained a Random Forest classifier on the data gathered from the Allen Cell Type Database. A total of 1145 cells of different sub-types (556 excitatory cells, 180 Sst, 142 Vip, 140 PValb, 127 Htr3a) for which linear properties (Input Resistance, Time constant and Resting potential), Firing properties (Gain, Threshold, Adaptation of spike upstroke, downstroke, threshold, instantaneous frequency and spike width at half height), and first spike descriptors (threshold, peak and trough potential, width at half height, height, upstroke, and downstroke) have been computed. After training the best Random Forest Classifier on 70% of the dataset (see methods), the cell class of the remaining portion of the dataset was predicted.

We first tried to identify the excitatory or inhibitory profile of the cell based on the genetic markers expressed by the cell (i.e.: Pvalb, Sst, Htr3a and Vip cells were considered as Inhibitory cells, the others as Excitatory cells). After hyper parameters grid search, a Random Forest classifier was able to efficiently differentiate between the excitatory and inhibitory cells from the Allen Cell Type Database, reaching Accuracy and F1-Score of 90.6% and 90.6% on average respectively. Notably, this classifier performed equally well for both excitatory and inhibitory cells (Table **??**).

We further tested the capacity of this classifier, this time making the distinction between the different sub inhibitory cell types, that is PValb, Sst, Htr3a+ expressing cells. Note that for this step we gather together Htr3a and Vip expressing cells, as Vip expressing interneurons constitute a sub type of Htr3a cell class (Tremblay et al., 2016). This classification yielded average classification results of 86.2% and 81.1% Accuracy and F1-score respectively. Between the different cell classes tested, the PValb expressing interneurons were the most accurately predicted cells, with 88.4% F1-score, followed by Htr3a+ (79.5%) and Sst (75.4%) (Table **??**).

Finally, we tried used the same approach to class neurons according to a more detailed classification scheme (i.e.: Excitatory, Pvalb, Sst, Htr3a and Vip). This analysis achieved mean prediction accuracy of 92.1%(Table **??**). These results were better than those of previous works relying on spike waveform transformation (Ghaderi et al., 2018; Rodríguez-Collado and Rueda, 2021). Furthermore, the mean accuracy was better than the accuracy achieved by Ophir et al. (2024) when using random forest (82.4%) and comparable when using more complex algorithms as LSPIN (91.6%). On the other hand, our analysis achieved 73.5% F1-score on average, which is lower than when using LSPIN model (87.7%). Yet, the fact that the F1-score achieved is slightly better than the F1-score obtained in Ophir et al. (2024) using random forest (72.6%), while relying on far less features (16 in the TACO pipeline, and 41 in Ophir et al. (2024)), extracted from single experimental protocol (long-step current clamp in the TACO pipeline, while Ophir et al. (2024) relied on long square, short square, ramp and noise inputs from the Allen Cell Type Database) suggests that our approach represents a convenient way to objectively characterize electrophysiological profiles of the main neuronal types. From this analysis, it appears that excitatory neurons are the most distinguishable neuronal class (F1-score : 93.3%) followed by PValb neurons (F1-score : 86.4%), Sst (75.7%), Vip (63.2%) and Htr3a neurons (49.2%). These results are in agreement with some previous works (Rodríguez-Collado and Rueda, 2021), and in opposition with others (Ghaderi et al., 2018). Such heterogeneity of qualitative assessment suggests that the quality of the prediction notably depends on the type of features used to train the predictive model.

**Figure 10:**
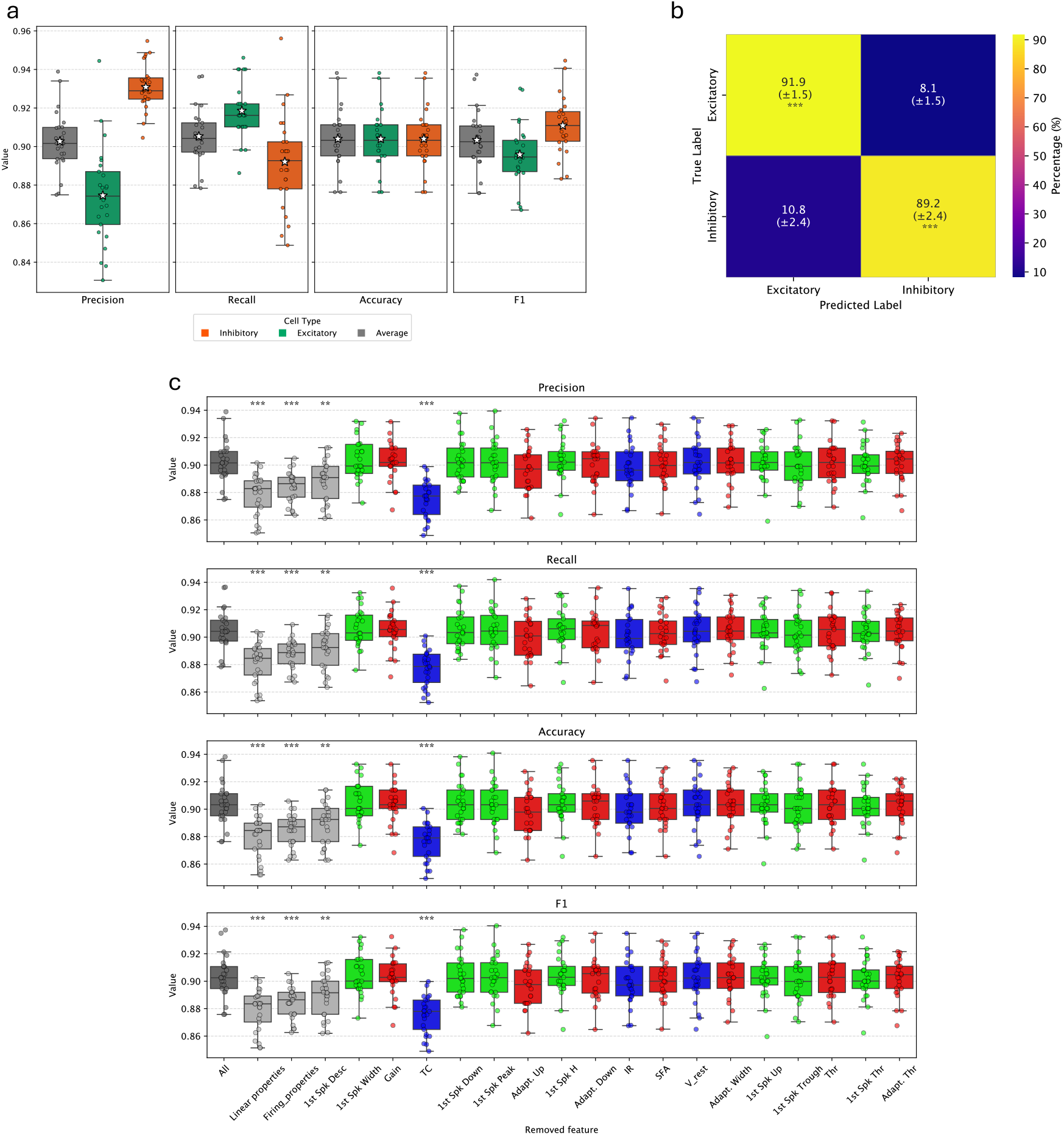
Classification performances of excitatory and inhibitory neurons from Allen Cell Type Database using random forest with grid search hyper parameters tuning over multiple iterations. **(a)** These plots represent multiple classification metric for the classification of excitatory and inhibitory neurons. Box plots represent the distribution of performances across iteration (individual points), with white stars indicating mean performance over iterations. **(b)** Classification confusion matrix for excitatory and inhibitory neurons (mean ± standard deviation). For each value, a t-test is performed to determine if the average value over iteration is significantly higher than chance level (here, 0.5). **(c)** Feature importance for excitatory and inhibitory neuron classification. The different panels represent the different classification metrics. For each metric, the “All” box plot represent the classification performance using all features (same as “Average” box plots from **(a)**), while other box plots represent performance with specific set of features removed (Linear or Firing properties or 1st spike descriptors), or single features removed, classified in descending order of feature importance when all features are used. Green, red and blue boxplots represent performance when single 1st spike descriptor feature, firing property or linear property is removed respectively. For each scenario, a t-test was performed to identify significant performance drops compared to classification with all features.

**Figure 11:**
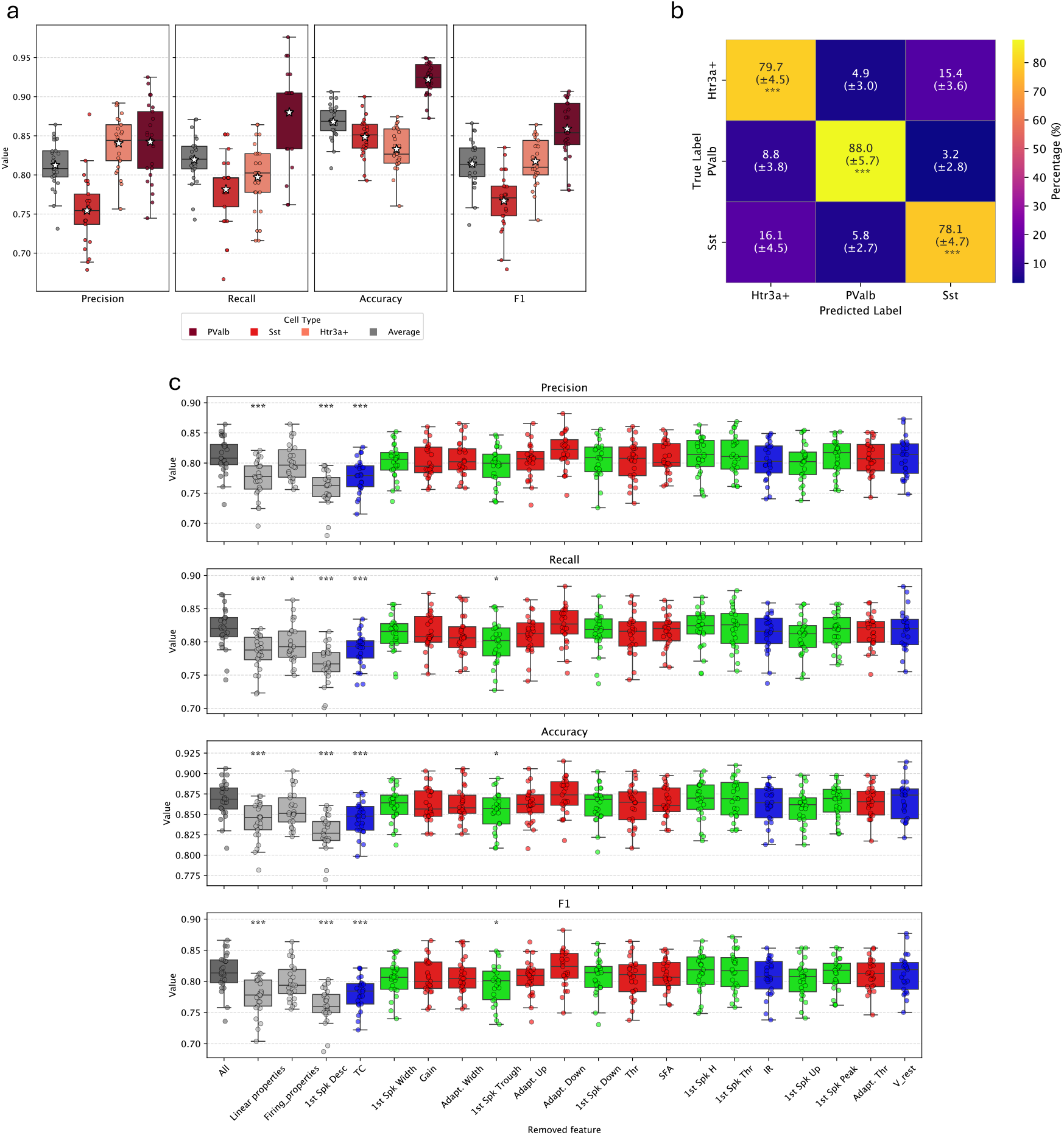
Classification performances of inhibitory sub-types neurons from Allen Cell Type Database using random forest with grid search hyper parameters tuning over multiple iterations. **(a)** These plots represent multiple classification metric for the classification of Sst, Pvalb and Htr3a+ neurons. Box plots represent the distribution of performances across iteration (individual points), with white stars indicating mean performance over iterations. **(b)** Classification confusion matrix for different inhibitory subtypes neurons (mean ± standard deviation). For each value, a t-test is performed to determine if the average value over iteration is significantly higher than chance level (here, 0.5). **(c)** Feature importance for inhibitory subtypes neuron classification. The different panels represent the different classification metrics. For each metric, the “All” box plot represent the classification performance using all features (same as “Average” box plots from **(a)**), while other box plots represent performance with specific set of features removed (Linear or Firing properties or 1st spike descriptors), or single features removed, classified in descending order of feature importance when all features are used. Green, red and blue boxplots represent performance when single 1st spike descriptor feature, firing property or linear property is removed respectively. For each scenario, a t-test was performed to identify significant performance drops compared to classification with all features.

#### 4.2.2 Leverage from a database to predict genetic markers in another database

We pursued our investigation by assessing the generalization of these features, in the context of inferring metadata from a database to another one. To do so, we used the random forest classifiers trained on the Allen Cell Type Database and applied these on cells obtained from the Scala 2019 database (Scala et al., 2019). Cell recorded in the visual cortex and genetically labeled (PValb (74), Sst (78), Vip (29), or Excitatory(37)) were classified first on an Excitatory or Inhibitory basis using the best random forest classifier trained on the Allen database (see Table **??**), then differentiating between the inhibitory sub cell times (see Table **??**), and finally between different excitatory neurons and the different inhibitory classes (Table **??**).

It resulted that a random forest model trained on the Allen CTD Excitatory and Inhibitory classes achieved a mean classification accuracy of 82.2% and a mean F1-score of 74.4% (Table 1) when predicting Excitatory and Inhibitory classes in cells from the Scala 2019 database. Yet the excitatory cells seemed less accurately predicted than inhibitory cells (F1-score 60.2% and 88.5% respectively)

**Table 1:** Performance of Scala 2019 VIS cells’ excitatory or inhibitory profile prediction using Random forest classifier trained on Allen Cell Type Database.

| Measure | Average | Excitatory | Inhibitory |
| --- | --- | --- | --- |
| **Precision** | 0.717 | 0.470 | 0.963 |
| **Recall** | 0.828 | 0.838 | 0.819 |
| **Accuracy** | 0.822 | 0.822 | 0.822 |
| **F1** | 0.744 | 0.602 | 0.885 |

These performances decreased when increasing the granularity of the classification (i.e.: increasing the number of cell type to predict), reaching an mean accuracy of 63.7% and a mean F1-score of 53.6% when considering only inhibitory cell types (Table 2), and 71.0% accuracy and 49.8% F1-score when classifying all different excitatory and inhibitory cell types (Table 3). Note that in the case where the model is being trained on a database containing Htr3a, it classified some of the Scala 2019 cells as such, whereas it doesn’t contain any, therefore resulting in precision = 0, Recall and F1-Score undefined. On the other hand the accuracy is not null as it takes into account all the cell correctly identified as not belonging to this class.

**Table 2:**
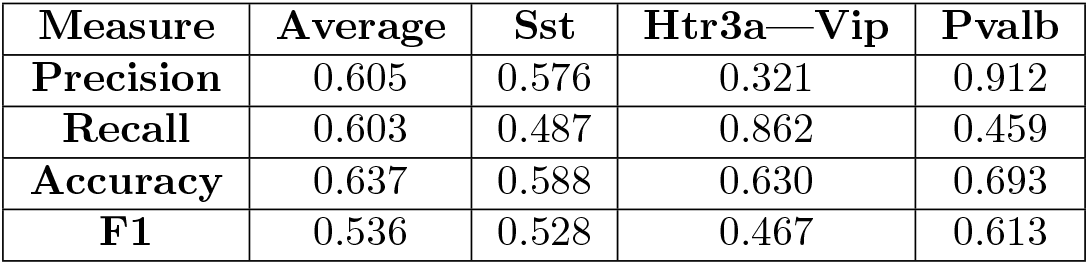
Performance of inhibitory cell type prediction of Scala 2019 cells using Random forest classifier trained on Allen Cell Type Database.

| Measure | Average | Sst | Htr3a—Vip | Pvalb |
| --- | --- | --- | --- | --- |
| **Precision** | 0.605 | 0.576 | 0.321 | 0.912 |
| **Recall** | 0.603 | 0.487 | 0.862 | 0.459 |
| **Accuracy** | 0.637 | 0.588 | 0.630 | 0.693 |
| **F1** | 0.536 | 0.528 | 0.467 | 0.613 |

**Table 3:**
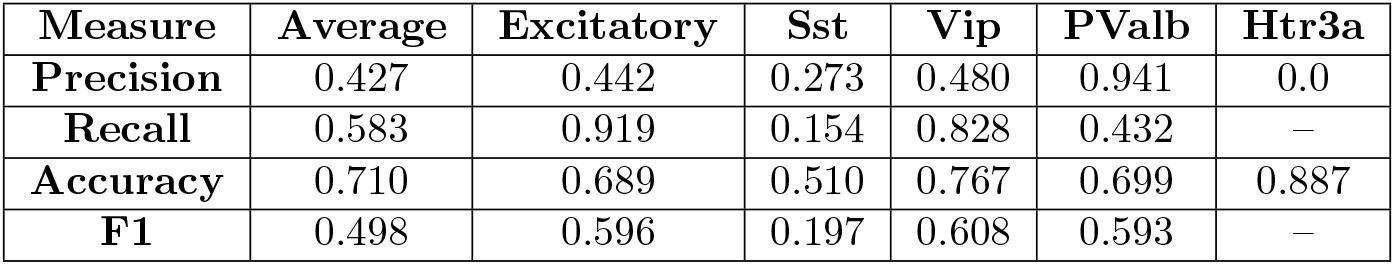
Performance of detailed cell type prediction of Scala 2019 cells using Random forest classifier trained on Allen Cell Type Database.

| Measure | Average | Excitatory | Sst | Vip | PValb | Htr3a |
| --- | --- | --- | --- | --- | --- | --- |
| **Precision** | 0.427 | 0.442 | 0.273 | 0.480 | 0.941 | 0.0 |
| **Recall** | 0.583 | 0.919 | 0.154 | 0.828 | 0.432 | — |
| **Accuracy** | 0.710 | 0.689 | 0.510 | 0.767 | 0.699 | 0.887 |
| **F1** | 0.498 | 0.596 | 0.197 | 0.608 | 0.593 | — |

We hypothesized that this decrease in cell class predictive performances could result from database-specific differences, in other word protocol-dependent parameters that could influence the extracted features and that the TACO pipeline cannot correct. For example, a noticeable difference arose from the Allen CTD and the Scala 2019 database, in that during the recording session, the Allen CTD used synaptic blockers in their recording solution, whereas the Scala 2019 did not (Gouwens et al., 2019; Scala et al., 2019). Such difference in the recording protocol could greatly affect the neurons’ biophysics, and thus the features extracted from the recordings. To confirm this hypothesis, we trained a Random Forest classifier on either Visual Allen CTD cells (using solely PValb, Sst and Vip cells to match to the Scala 2019 database), Visual Scala 2019 database, or on both databases, and tried to predict a second subset of cells from the Scala 2019 database recorded in the Somato-sensory cortex using the same experimental procedure, and targeting only Sst neurons (69 cells). It appears that when using solely a subset of the Allen CTD to train the classifier, classification of Sst cells recorded in the Scala 2019 database achieved poor performance by identifying less than half of the cells as Sst. These performances increased when using a combination of both Allen CTD and Scala 2019 database to train the random forest, and even slightly better when solely using Scala 2019 to train the model (Table 4). These results illustrate the fact that the training dataset has a great impact on the predictive performance for an external database. Indeed, a training database, even sampled similarly, may achieved worse results (in this case half worse) than another one. The reason explaining such differences may arise from the difference in experimental conditions and protocols impacting the absolute values of the features, and therefore preventing efficient classification. Although it should be noted that a classifier trained on both databases achieved almost as good results as a classifier trained using data from the same database. This suggests that to achieve better performance of neuronal type classification, one can benefit from a collection of different database to encompass the features differences resulting from differences in experimental procedures.

**Figure 12:**
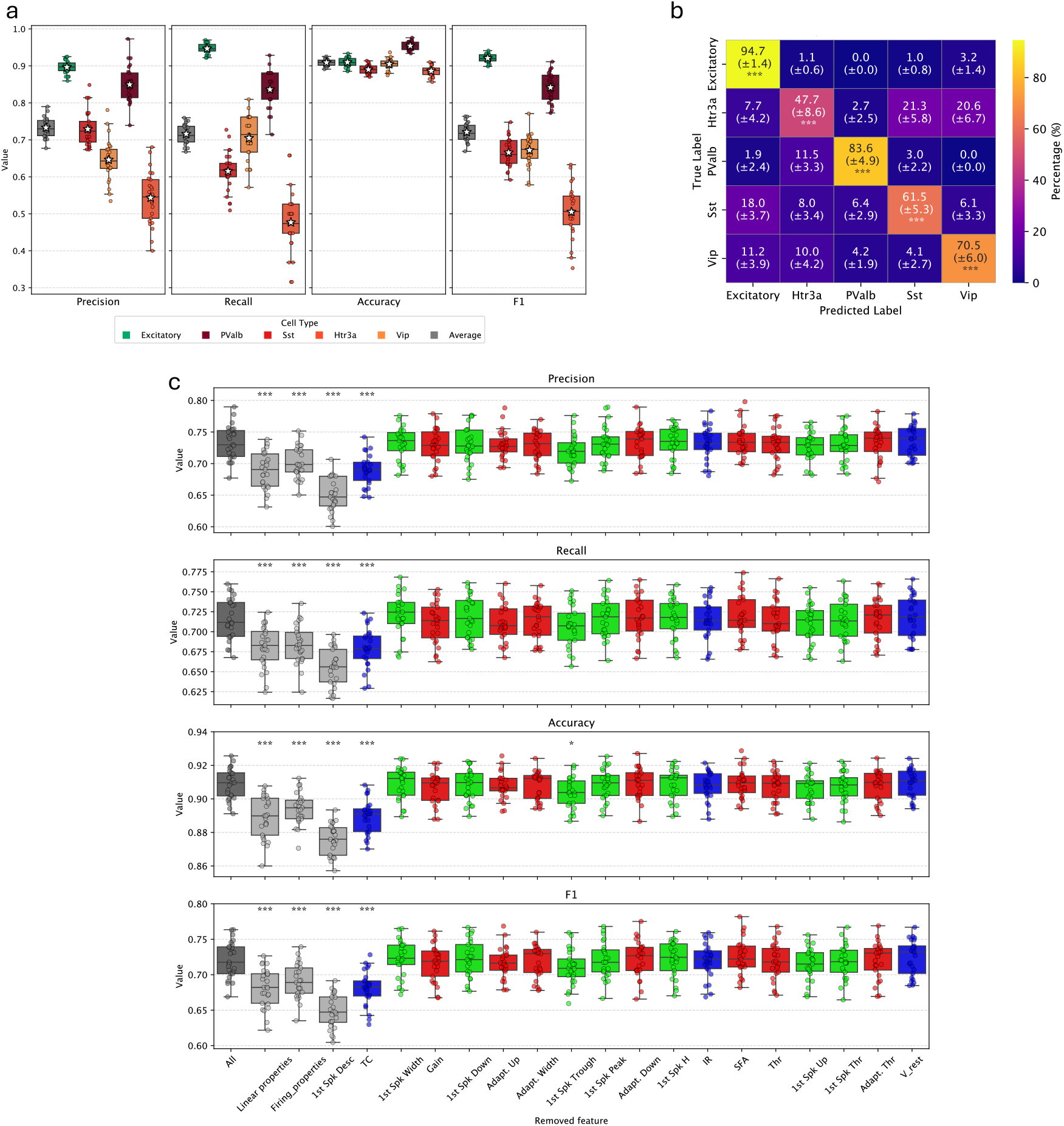
Classification performances of different neuronal types from Allen Cell Type Database using random forest with grid search hyper parameters tuning over multiple iterations. **(a)** These plots represent multiple classification metric for the classification of Excitatory and inhibitory Sst, Pvalb, Vip and Htr3a neurons. Box plots represent the distribution of performances across iteration (individual points), with white stars indicating mean performance over iterations. **(b)** Classification confusion matrix for different neuronal types (mean ± standard deviation). For each value, a t-test is performed to determine if the average value over iteration is significantly higher than chance level (here, 0.5). **(c)** Feature importance for neuronal types classification. The different panels represent the different classification metrics. For each metric, the “All” box plot represent the classification performance using all features (same as “Average” box plots from **(a)**), while other box plots represent performance with specific set of features removed (Linear or Firing properties or 1st spike descriptors), or single features removed, classified in descending order of feature importance when all features are used. Green, red and blue box plots represent performance when single 1st spike descriptor feature, firing property or linear property is removed respectively. For each scenario, a t-test was performed to identify significant performance drops compared to classification with all features.

**Table 4:** Performance of Sst cell type prediction of somatosensory Scala 2019 cells using Random forest classifier trained on either Allen Cell Type Database (using only Vip, Sst, PValb), Visual cortex cell Scala 2019, or both. The column with cell types indicate the number of cell predicted as such, knowing that the subset of cells to class contained 69 cells.

| Train Database | Sst | PValb | Vip | Accuracy | F1-Score | Recall | Precision |
| --- | --- | --- | --- | --- | --- | --- | --- |
| **VIS Allen CTD** | 27 | 19 | 23 | 0.391 | 0.563 | 0.391 | 1.0 |
| **VIS Scala 2019** | 65 | 4 | 0 | 0.942 | 0.970 | 0.942 | 1.0 |
| **Both** | 53 | 14 | 2 | 0.768 | 0.869 | 0.768 | 1.0 |

## 5 Conclusion and Perspectives

The implementation of general methods for the quantification of neuronal biophysical properties represent a major aspect of computational neuroscience. While the definition of such properties are commonly shared (e.g., gain, rheobase…), their reliable and coherent measurement from diverse current-clamp protocol experiments is a challenge. We introduced new methods to characterize neuronal I/O relationship, and measure functional properties such as gain, rheobase, saturation and SFA in a way which is robust to the differences in experimental protocols. These methods provided measures which lead to better classification performance of neurons according to their neuronal type, compared to previous works relying on other methods. These methods were designed to be applied on different databases of current-clamp recordings despite the differences in experimental protocol, therefore enabling cross-database analysis of neuronal functional properties.

Another approach of inter-database comparison is the NeuroElectro database (Tripathy et al., 2014, 2015), which, based on a text-mining algorithm, has gathered values for multiple properties in various parts of the brain. This work represented an unprecedented valorization of older electrophysiological studies. The framework consisted of identifying relevant articles by massively downloading articles from neuroscience journals and selecting those containing keywords like “*input resistance*” or “*resting membrane potential*,” focusing on articles likely to contain relevant information to gather. Their approach, mixed between a semi-automatic extraction pipeline and a human-based curation allowed them to gather more than 2300 electrophysiological measurements from 335 references, representing a major opportunity to leverage existing works. However, as stated by the authors, it supposes that a given electrophysiological property has been measured similarly by the different authors, notably using similar protocols, then that the properties was computed in the same way. Finally, for the databases to be comparable, the experimental conditions should be similar, and the data acquired similarly. To overcome this, the reported data were, when possible, normalized to common reference definitions, otherwise not considered. However, this was done manually, based on the method described in the article, and therefore could add unexplained and unrealistic variance to the global data, without ways of identifying it. This is notably true for diverse experimental artifacts that may affect numerous aspects of reported neuron electrophysiology like bridge error. Also, as the data reported in articles, need to be constrained and readable, they are often presented in the mean±standard deviation format, which represent an important source of data reduction, and simplification(Speelman and McGann, 2013).

Other initiatives, as the establishment of common features definition and data structures have also been made to improce interoperability of neurosciences data. For example, the Petilla convention aimed at providing general definitions for multiple aspects of cortical GABAergic neurons to facilitate their common classification (Ascoli et al., 2008). Regarding the data structure in the neuroscience community, the Neurodata Without Border (NWB) data format has been proposed as a common language for the sharing and interoperability of neurophysiological data (Teeters et al., 2015). The goal is notably to propose a single neurophysiology framework for the diverse types of protocols commonly used in neurosciences, like intra-cranial electrophysiology probes, electrocorticography, or intracellular recording of membrane potentials, and their analysis and visualization. It has been designed to co-evolve with neurophysiological research. Indeed, by being open source, it enables different teams to modify it, and adapt it to their particular research (Rübel et al., 2022). A particular example of an intracellular electrophysiological database published in the NWB format is the Allen Cell Type Database (Gouwens et al., 2019), making it an unskippable and reusable reference for other studies (Ghaderi et al., 2018; Nandi et al., 2022).

These initiatives illustrate the need for computational neuroscience to embrace more generally the Open Science practices to ensure higher interoperability of databases composed of similar data. Indeed, the cross compatibility between multiple experimental databases represents a key aspect of modern neurosciences, particularly with respect to the open sciences principles and the need to reduce animal experiment (Janssens et al., 2023; Richter, 2024). While research projects naturally aim at investing particular questions and so require specific experimental paradigm and conditions, we believe some of the publicly available data can be of interest to other labs. Furthermore, as we saw, different experimental paradigms and quantification methods can lead to erroneous interpretation of comparison between different research projects. While some projects have targeted this question of inter compatibility of neurophysiological data by proposing common data format (Rübel et al., 2022), the vast majority of publicly available database adopt lab-based database organization and file format, making the cross analysis of neurophysiological data complicated.

To overcome this observation, we proposed a data structure independent python-based data analysis pipeline, with detailed, data-based and robust quantification methods to enable coherent databases integration. The TACO pipeline has been designed to enable any lab with a database of intracellular recordings to analyze and coherently compare its database with others’, relying only on basic descriptive information and on small database-specific python code for data extraction. To ensure coherent comparison of data different aspects, either experimental like the bridge error estimation, or physiological like Type I and Type II neurons distinction are addressed. Also, the user can input custom quality criteria to accept or reject individual traces, before performing analysis, to ensure applying same quality criteria to all databases. Along these different considerations to ensure data quality, the different analysis are detailed, rely on preliminary estimations based directly on the data, and can be visually inspected using the cell data application. Furthermore, the careful characterization of the electrophysiological profile of neurons based on long-step current-clamp recording is a major aspect of the pipeline, and constitute an important step toward better valorization of intra-cellular recording database. Indeed, it provides a consistent framework to predict cellular classes of neurons based on a limited number of features (i.e.: linear properties, firing properties and first spike descriptors), and reaching higher prediction performances than similar previous works, and equally acceptable performance than more sophisticated machine learning methods relying on richer data (i.e.: obtained from multiple different recording protocols). Yet it appears that to infer this kind of information between different databases, training a model on a aggregated database recorded under different lab-specific protocols is needed, as it tends to reduce the model over-fitting to a particular dataset.

Proposing a common data analysis pipeline, supporting methods suitable for every protocol of current-clamp experiments, that can be applied on any databases, regardless of its original structure, and enabling common and custom quality assessments, represent an important valorization perspective of publicly available databases. We believe that this common data-analysis approach is complementary with the need of a common data structure like NWB which greatly enhance the cross-databases analysis. In the future, on the same model than our framework to enable the general application of user-defined quality criteria, the pipeline could be upgraded with the possibility of performing user-defined analysis, while still limiting the requirement for in depth-modification of the pipeline.

## 6 Materials and methods

### 6.1 Software packages

The pipeline has been implemented using Python 3.11, distributed via Anaconda. The cell viewer application has been designed using the Shiny for Python framework. The fit of the pipeline are done using the lmfit library (Newville et al., 2015). The minimization of the objective function was set to default (i.e.: Levenberg-Marquart) for all fits, except for the Hill-Sigmoid fit to original data (Section 3.4.3), where the Least-Squares minimization using Trust-Region Reflective method was also tested. Whatever methods yields best results (i.e.: lowest fit error) was used for a given cell.

The statistical analysis were performed in R programming language (v 4.3.1), using rstatix package.

### 6.2 Statistical analysis

Before performing the different statistical analysis of mean differences, the parametric hypothesis were tested to decide which test to use. To test if the data were normally distributed per group, we used Shapiro-Wilk test, while the Levene’s test was used to assess the homogeneity of the variance. The observations were independent from each others. If either of the two preliminary tests was statistically significant (*p <* 0.05), then the variance test applied was the non-parametric Kruskal-Wallis to assess if there were any statistically significant difference between means of the groups of interest. If so (i.e.: *p <* 0.05) then a group-wise comparison test was performed using the Dunn’s test, with Bonferroni adjusted p-values. In the cases where both Shapiro-Wilk and Levene’s tests were not statistically significant (i.e.: *p* ≥ 0.05), then a one-way ANOVA test was used to assess any statistically significant difference between means of the groups of interest. If so, then post-hoc Tukey test was used for pair-wise comparisons, with p-values corrected using Bonferroni methods. The p-values reported on the plots represent the following levels of statistical differences : Not significant : *p <* 0.05, * : 0.05 ≤ *p <* 0.01, ** : 0.01 ≤ *p <* 0.001, * * * : 0.001 ≤ *p <* 0.0001, * * ** : *p* ≤ 0.0001

### 6.3 Random forest classification

#### 6.3.1 General design

To test the classification performances of the different features extracted as part of the TACO pipeline, we implemented a repeated train-test split and cross validation framework for random forest classifier. A set of 17 features of different nature were extracted. Linear properties: Time constant (ms), input resistance (GΩ), and resting potential (mV); Firing properties : gain (Hz/pA), threshold (pA), Adaptation of instantaneous frequency, spike width, spike threshold, upstroke and downstroke (Firing properties); 1st spike descriptors: Threshold potential (mV), height (mV), width (ms), upstroke (mV/ms), downstroke (mV/ms), peak potential (mV) and trough (mV).

To mitigate the influence of data splitting variability, the following procedure was repeated 25 times using different random seed for each iteration. For each iteration, the dataset was randomly split into train (70%) and test (30%) datasets. Hyperparameters optimization was performed on the training dataset, using grid search strategy to select the combination of maximum depth (tested between 5 and 20 with a step of 2), minimum sample split (tested between 10 and 25 with a step of 2) and maximum leaf nodes (tested between 20 and 26 with a step of 2), leading to the best performances. Gini impurity was used as the splitting criterion. The optimization process aimed to maximize the weighted F1-score, which accounts for class imbalances by weighting the F1-score according to the number of instances in each class. The random forest with the hyperparameter combination leading to the best classification performances over a 10-fold cross validation on the training dataset, was selected, and its performances evaluated on the held-out test dataset.

To test classification performance across database, the random forest was trained on a first database as described above, and the best resulting classifier was then used to predict data from another database.

#### 6.3.2 Classification performances evaluation

To evaluate the performance of our classification model, we computed several standard metrics, including precision, Recall, Accuracy, and F1-score. These metrics provide a comprehensive understanding of the classifier’s performance, particularly in the context of class imbalance, which was a key consideration in this study.

- **Precision** is the proportion of true positive predictions out of all instances classified as positive by the model. It is calculated as:

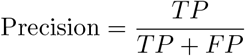

where *TP* refers to the true positives and *FP* refers to the false positives. Precision indicates the model’s ability to avoid misclassifying negative instances as positive (i.e.: False positive).
- **Recall** (also known as Sensitivity or True Positive Rate) is the proportion of true positive instances out of all actual positive instances in the dataset. It is calculated as:

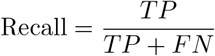

where *FN* refers to the false negatives. Recall reflects the model’s ability to capture all relevant positive instances.
- **Accuracy** measures the proportion of correctly predicted instances (both positive and negative) out of the total number of instances. It is calculated as:

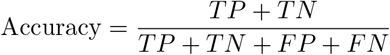

where *TN* refers to the true negatives. Accuracy is a general measure of the model’s correctness but may be less informative in the presence of class imbalances.
- **F1-score** is the harmonic mean of Precision and Recall, offering a single metric that balances both concerns. It is computed as:

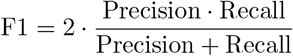

The F1-score provides a more balanced metric when there is a trade-off between Precision and Recall, and it is especially useful in contexts where false negatives and false positives have similar costs.

## 7 Supplementary figures

**Figure 13:**
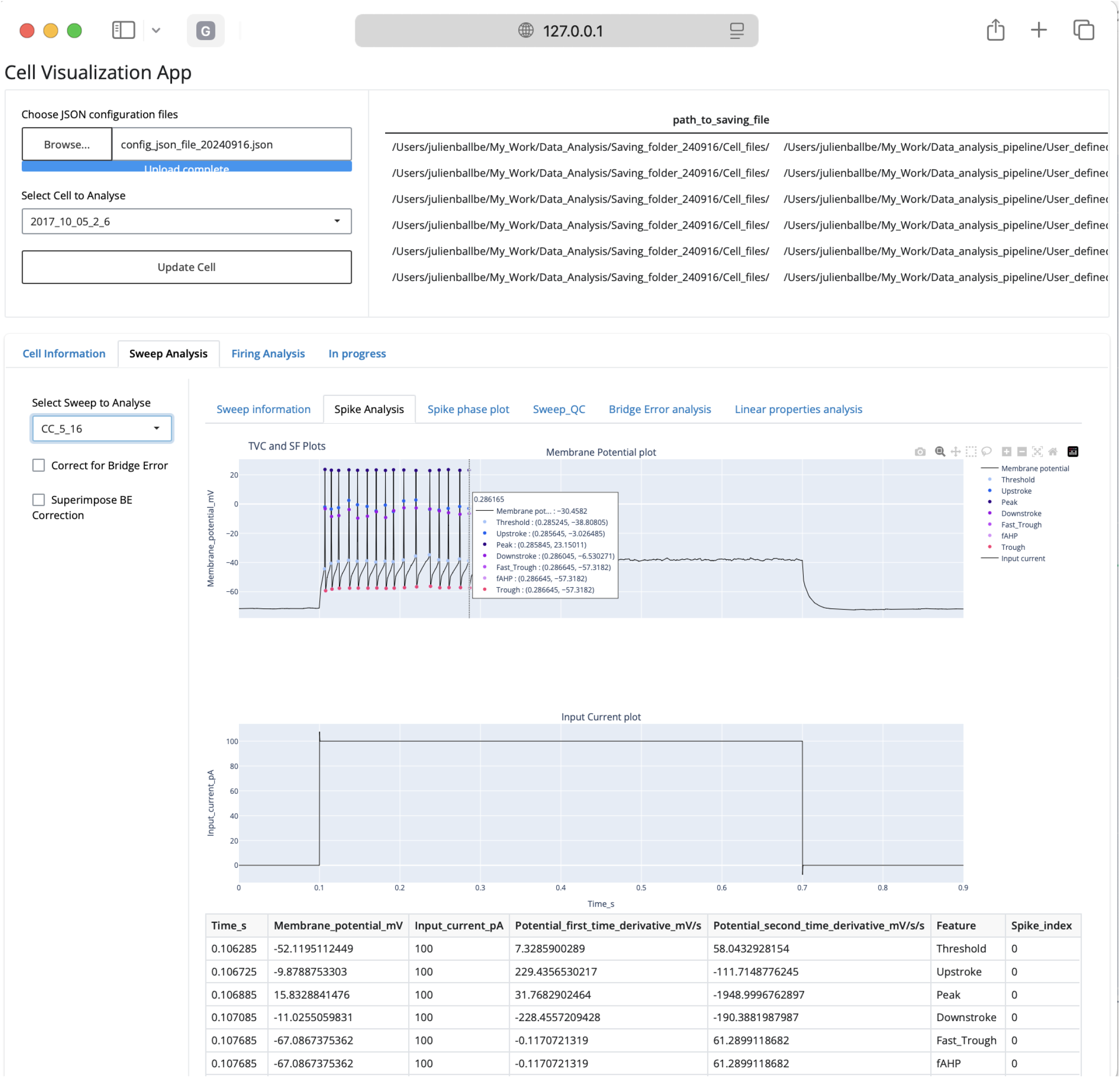
Cell Visualization App, Spike analysis panel.

**Figure 14:**
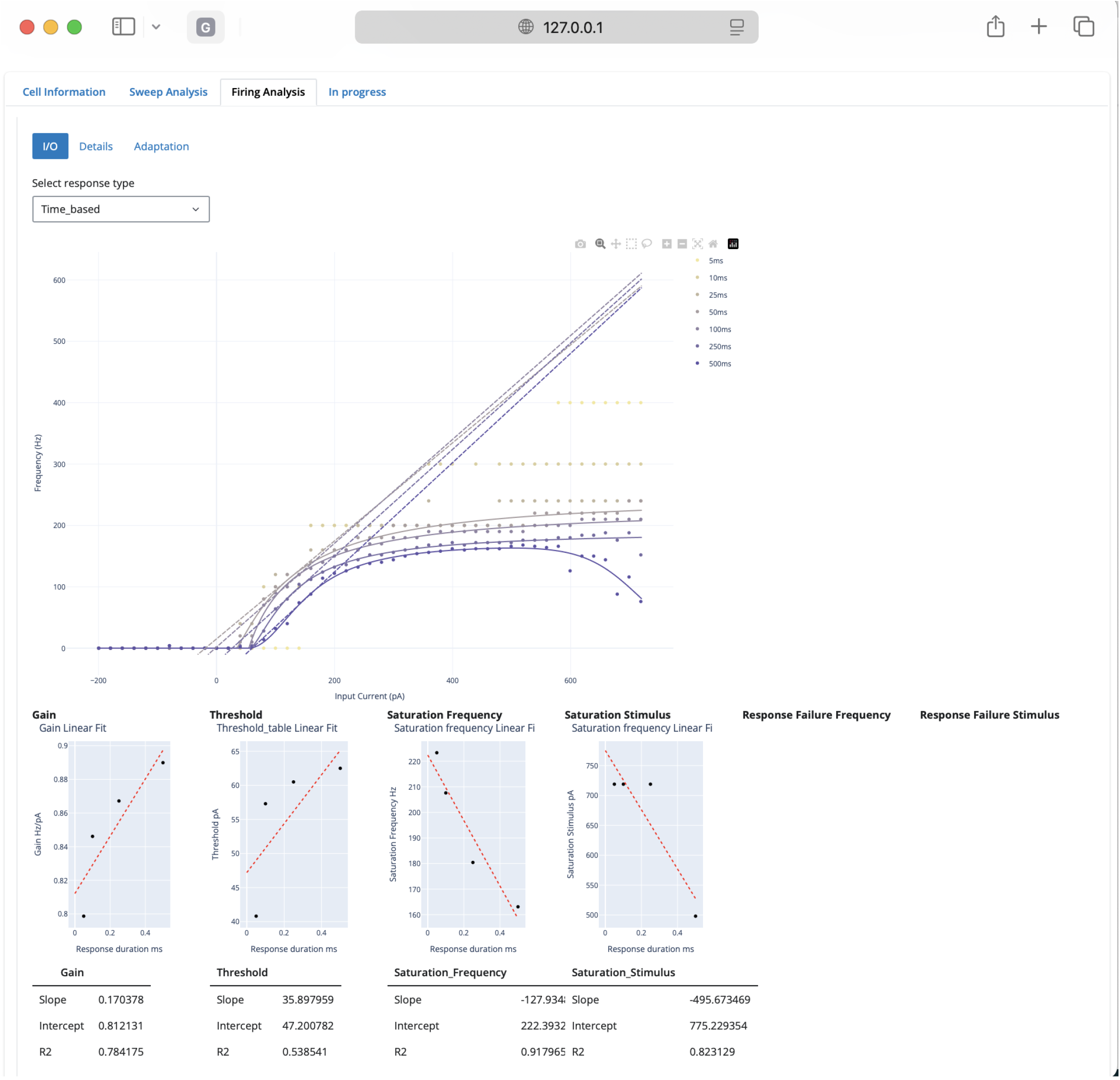
Cell Visualization App, I/O fit panel.

**Figure 15:**
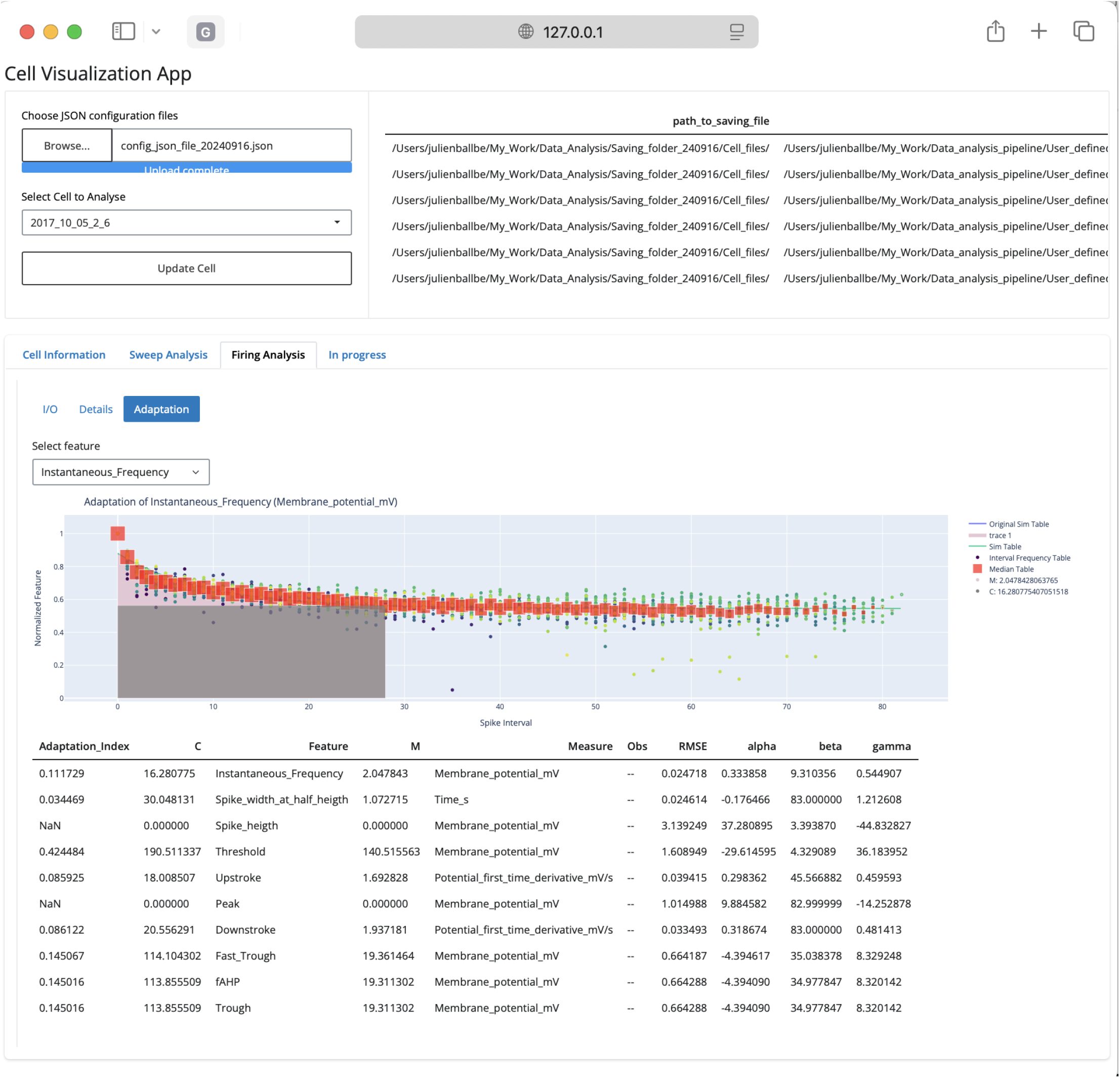
Cell Visualization App, Adaptation panel.

**Figure 16:**
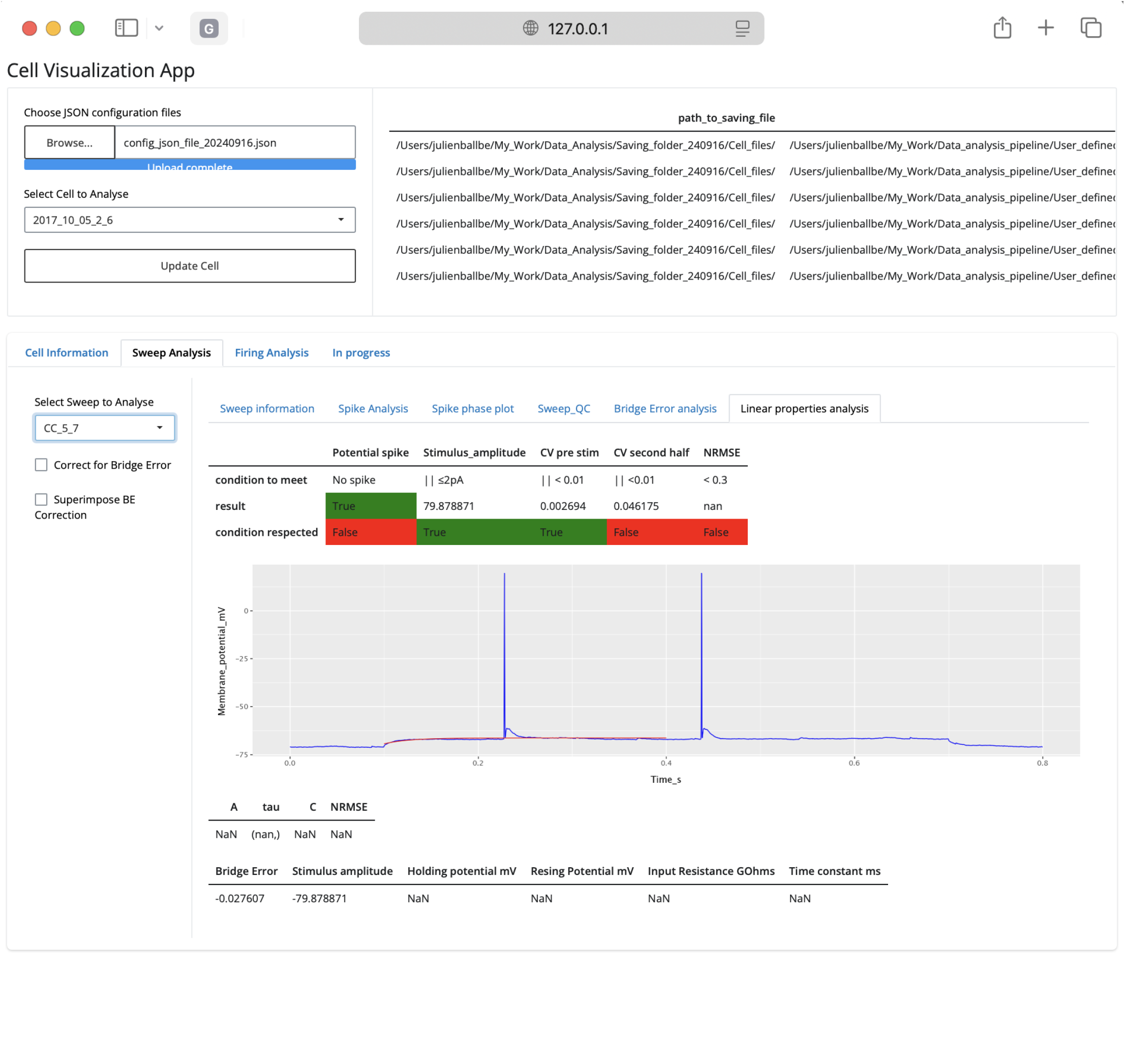
Cell Visualization App, Linear properties panel.

**Figure 17:**
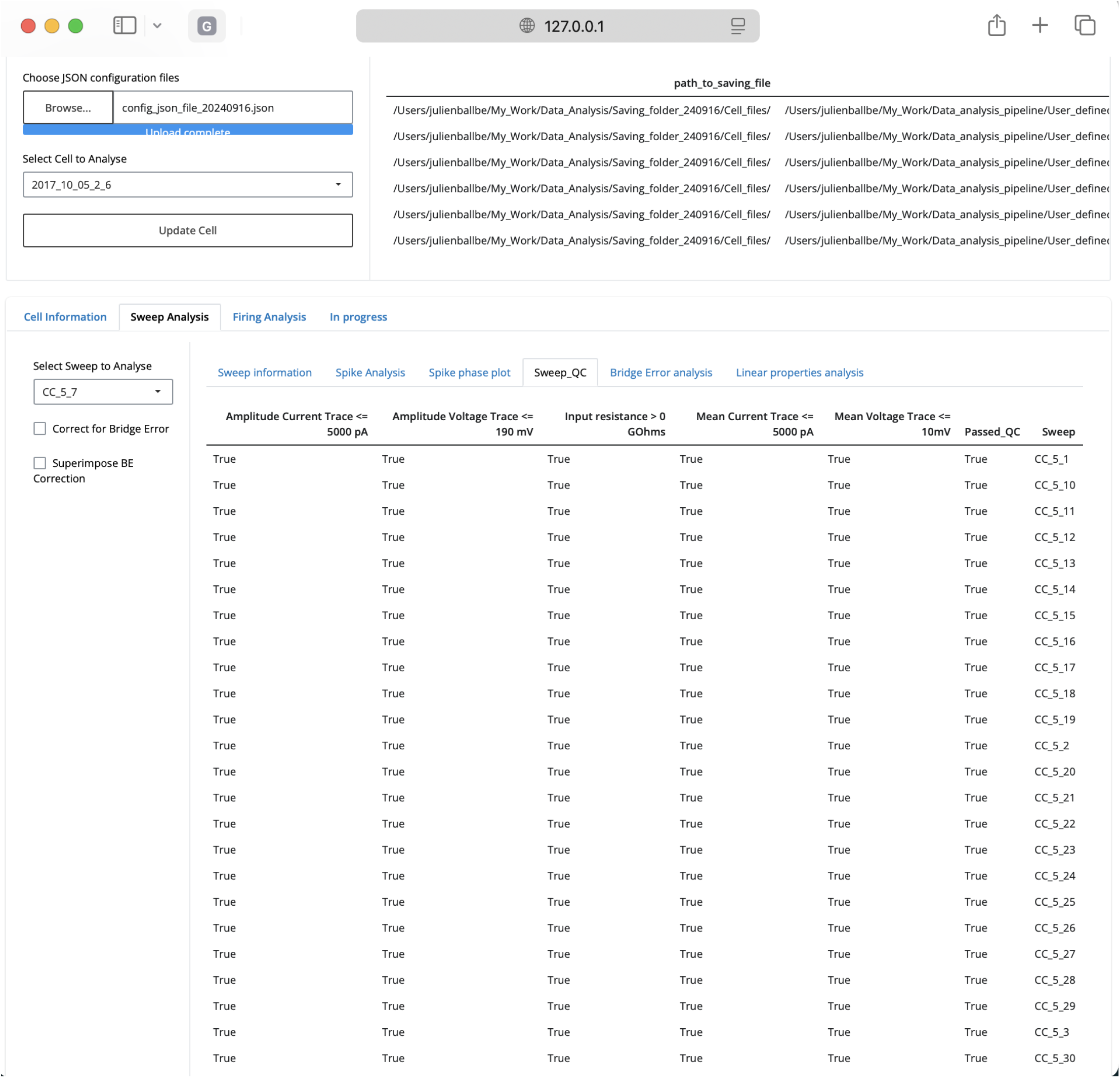
Cell Visualization App, Sweep QC panel.

